# Approximate Bayesian computational methods to estimate the strength of divergent selection in population genomics models

**DOI:** 10.1101/2023.06.06.543823

**Authors:** Martyna Lukaszewicz, Ousseini Issaka Salia, Paul A. Hohenlohe, Erkan O. Buzbas

## Abstract

Statistical estimation of parameters in large models of evolutionary processes using SNP data is often too computationally inefficient to pursue using exact model likelihoods. Approximate Bayesian Computation (ABC) to perform statistical inference about parameters of large models takes the advantage of simulations to bypass direct evaluation of model likelihoods. We use forward-in-time simulations of a mechanistic model of divergent selection with variable migration rates, modes of reproduction (sexual, asexual), length and number of migration-selection cycles, and investigate the computational feasibility of ABC to perform statistical inference and study the quality of estimates on the position of loci under selection and the strength of selection. We evaluate usefulness of summary statistics well-known to capture the strength of selection, and assess their informativeness under divergent selection. We also evaluate the effect of genetic drift with respect to an idealized deterministic model with single-locus selection. We discuss the role of the recombination rate as a confounding factor in estimating the strength of divergent selection, and we answer the question for which part of the parameter space of the model we recover strong signal for estimating the selection and make recommendations which summary statistics perform well in estimating selection.

## 1 Introduction

Divergent selection occurs when populations adapt to contrasting environments, causing the accumulation of genomic differences due to differential selective pressure in these environments. Identity of loci under divergent selection and estimating the strength of divergent selection at these loci play a key role in detecting divergent selection, which can be a driving force of speciation (Butlin et al. 2012). Here, we aim to build theoretical and experimental models of divergent selection and assess the computational feasibility and quality of statistical inference under these models. Specifically we investigate whether the loci under divergent selection can be identified, and the strength of selection can be estimated with reasonable precision using state of the art simulation-based statistical methods.

Our experimental system is the baker’s yeast (*Saccharomyces cerevisiae*) as a fast-evolving model organism. Yeast is an eukaryotic model organism with a moderate number of linear chromosomes (n=16) and a genome-wide number of cross-over events per meiosis that is comparable to larger eukaryotes (Mancera et al. 2008), but the small genome size (12 Mb). Yeast can undergo both asexual and sexual reproduction. Crossing of yeast strains and meiosis (sporulation) can be experimentally controlled. The sporulation when sexual reproduction takes place requires starvation of yeast from nitrogen, glucose, and carbon source (Mitchell 1994). Single haploid spores can be isolated and sequenced to determine haplotype phase across the genome as well as precisely map meiotic crossover, gene conversion events, and recombination rate heterogeneity across the genome (Mancera et al. 2008).

Some of the challenges in detecting divergent selection are as follows. On the statistical side, divergent selection models that we investigate are the result of stochastic processes on genomes through generations and therefore, they are large models. This causes two main challenges for statistical inference. The first is formulating a workable exact likelihood function. Simulation-based computational methods working with approximate model likelihoods as opposed to exact likelihoods partially solves this problem. The second is making inference scalable for large simulation studies to investigate model properties so that we know what to expect when we perform inference using data. A well-known simulation-based method based on approximate likelihoods is Approximate Bayesian Computation (ABC). ABC uses simulated data under an assumed model using a large set of parameter values to generate an approximate sample from the posterior distribution of interest by bypassing the evaluation of the likelihood function (Beaumont et al. 2002).

On the genetics side, physical linkage of nearby neutral loci to the locus under selection may lead to genetic hitchhiking (Barton 2000) (i.e., the change in allele frequency at neutral loci), which contributes to the formation of genomic islands (Nosil & Feder 2012), thereby making identifying the locus under selection challenging. The magnitude of the hitchhiking effect is modeled to be proportional to the distance of the actual locus under selection, influenced by the recombination rate (Qanbari 2020), for which we apply an average, fixed genomic recombination rate.

Keeping these two main difficulties in perspective, we design a computationally efficient simulator scalable at genomic scales to study the behavior of a divergent selection model. We use our simulator in a large simulation study to assess which statistics are informative about divergent selection, under variable migration, mode of reproduction (sexual or asexual), and in the presence of fixed recombination. We also investigate the bias and variance of estimators in estimating the strength of selection.

To address these, in section 2, we describe the experimental design of the biological yeast system and model parameters controlled in laboratory settings, the developed theoretical population genetics model, and how the experimental design of the theoretical model translates to the biological model. Next, in section 3, we describe the constraints of the process of generation of the data, define the model parameters, and we build the simulator for the theoretical population genetics model. In section 4 we describe the ABC, and how it assesses the summary statistics from the simulator output data in the estimation of the model parameters, with and without the initial outlier scan on the observed data where the outlier scan reduces the parameter space of potential positions under selection to be considered. In section 5 we present our results on estimating model parameters from both with and without the loci outlier scan, and for four different scenarios under which the initial parameters vary for the outlier scan method. To conclude, in section 6 we discuss the application of the simulation study combined with the ABC to bypass the likelihood function, the role of the parameter space of selection and for which part we recover a strong signal, what summary statistics out of the tested with our model outperform the others, and finally the confounding role of the recombination rate to make inferences about the biological yeast system.

## 2 Model

In this section, we first describe the biological system in which *yeast* is used as a model organism in an experimentally controlled environment of selective pressures, type of reproduction, and strength of migration. Then we describe a theoretical population genetics model as an idealization of this biological system.

### 2.1 Description of the biological system and experimental design

We have genetically engineered an obligate diploid cross between our two focal yeast strains, a North American oak isolate (YPS128) and an Australian vineyard isolate (DBVPG1106). The YPS128 and DBVPG1106 strains form the biological system experimental setup and they differ by over 70,000 SNPs, an average of 1 SNP per *∼*165bp (Hether 2016). We treat each SNP as a *locus*. The two strains form a diploid pool, without recombination, followed by sporulation of randomly selected two parents, during which recombination occurs, to create an offspring ancestral pool at time *t* = 0. Half of the ancestral becomes a founding population assigned to evolve in sodium dodecyl sulfate (SDS) and the other half of the ancestral pool becomes a founding population assigned to evolve in sodium chloride (NaCl). These environments induce differential selective environmental pressures in two populations.

The biological experiment was performed under four different selection-migration cycles treatments, with three replicates per treatment. the four different treatments were as follows: no migration and no sporulation (asexual reproduction), no migration and sporulation (sexual reproduction), 20% migration and sporulation, and 50% migration and sporulation. During the divergence with gene flow, the populations evolve asexually in one of the mediums for 5 days, which is assumed to be equivalent to *t^∗^* = 50 generations. Then, one of the four scenarios of migration conditional on the type of sexual reproduction is implemented for one generation. Thi cycle is repeated four times, resulting in evolution under divergent selection for approximately 200 generations.

### 2.2 Description of the theoretical population genetics model

We assume diploid organisms that differ by *L* bi-allelic loci. Each population is of constant effective population size *N_e_*, and they evolve in non-overlapping generations. We obtain the populations ***X*_0_** and ***Y*_0_** at time *t* = 0 by crossing the founding populations at all loci consisting of all private alleles at *L* loci, then incorporating the recombination events into the process, and finally we evolve the populations through generations as follows.

We assume that the two parental genomes for an offspring are uniformly randomly (independently of each other) distributed, one parent from each population. The number of recombinations on the offspring’s genome, *n_r_*, is binomially distributed with probability *r*. We define the position vector on which these *n_r_*events happen by ***L_r_***, ∥***L_r_***∥ ≤ *L*, at which recombination events are uniformly randomly (independently of each other) distributed on *L* loci. The two parental genomes are recombined in positions defined by ***L_r_*** to obtain the offspring genome.

The reproduction in recurring cycles of *t^∗^* generations: We start from *t* = 0, isolate ***X_t_*** and ***Y_t_*** from each other and implement asexual reproduction for a sequence of *t^∗^ −* 1 generations. These *t^∗^ −* 1 generations allow for population and loci-specific divergent selection to act on each population.

We let *s_i_* to denote the selection coefficient at locus *i ∈* {1, 2, . . ., ∥***L_s_***∥}, where the fitness of the reference allele *a_i_* under selection is (1 + *s_i_*) if population carrier has *a_i_* copy, else (1 + 0). We assume that selection effects are additive across loci on the genome such that the fitness of an individual at time *t −* 1, in population *j* (*j ∈* {***X***, ***Y*** }) (**Part A** of **Fig. 1**) is

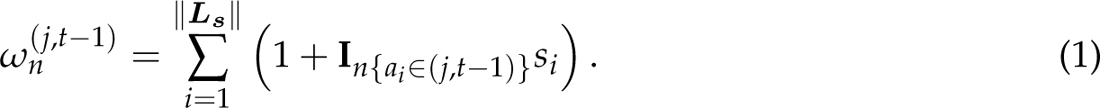

**Fig. 1.**
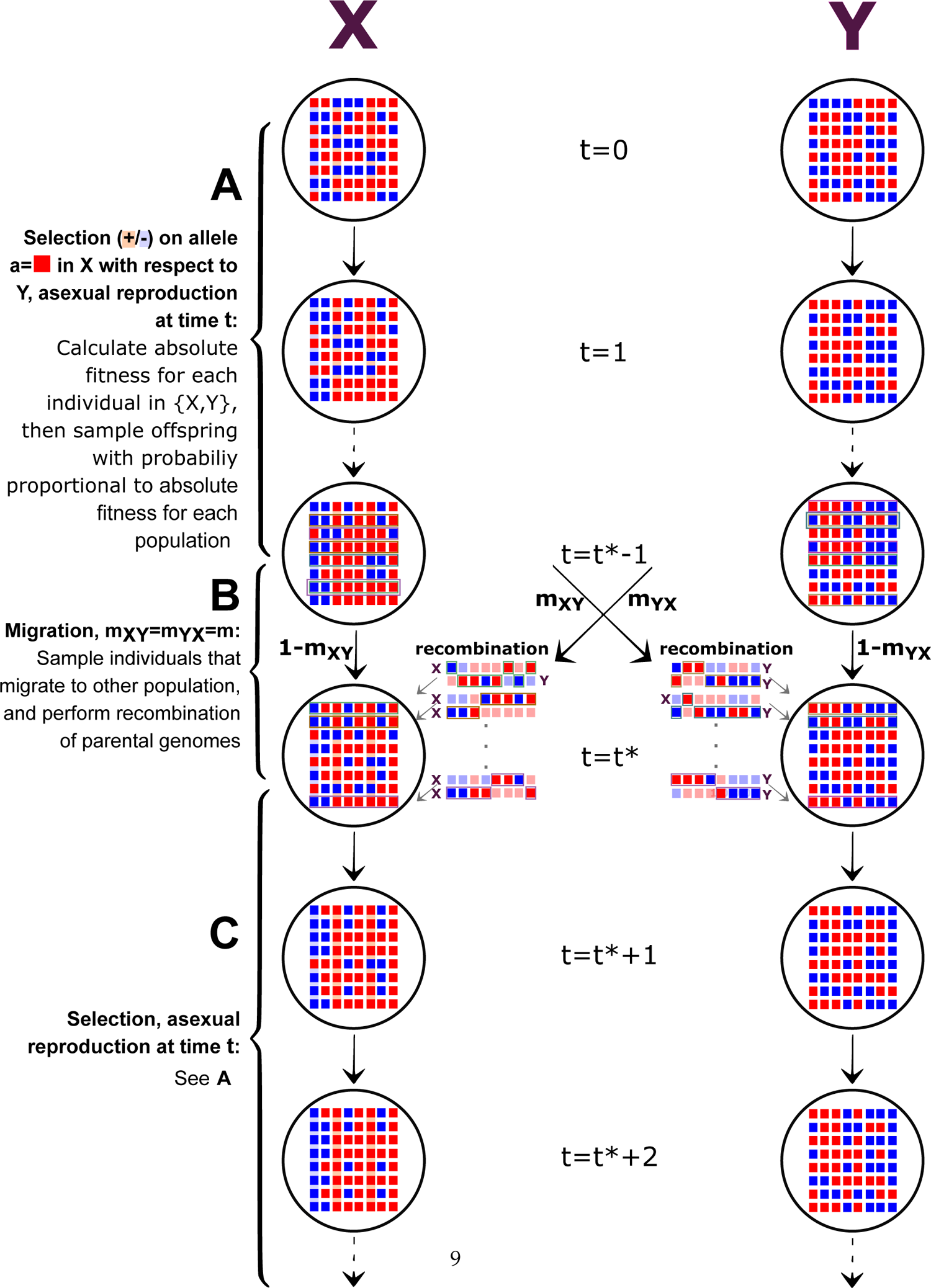
Population genetic model for divergent selection: A. Population divergence for *t^∗^* 1 generations during which reproduction is asexual and the absolute fitness depends on the allele-specific copy under which selection acts upon. Each offspring is genetically identical to its only parent and an individual is chosen to be a parent with probability proportional to its fitness. B. Symmetric migration rates *m_XY_* and *m_YX_* between *t^∗^ −* 1 and *t^∗^* generation. Neutral evolution with recombination. C. Second population divergence.

Then, for each distinct genome, the probability of including an offspring at generation *t* is multi-nomially distributed with the probability of successes proportional to their normalized fitnesses given by

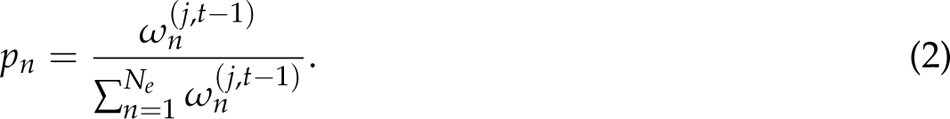

From generation *t^∗^ −* 1 to *t^∗^* populations undergo asexual or sexual reproduction with recombination and symmetric migration. Migration rate from ***X_t_∗_−_*_1_** to ***Y_t_∗*** and from ***Y_t_∗_−_*_1_** to ***X_t_∗*** is denoted by *m*. We sample uniformly randomly (independently of each other) *N_e_m* parents to migrate from population *j* to the other population. After migration, reproduction is either sexual (*sex* = 1), or asexual (*sex* = 0). If *sex* = 0, an offspring is an exact copy of a single parent chosen uniformly randomly. If *sex* = 1, we choose two parents uniformly randomly (independently of each other) from the same population. The recombination steps are the same as described above (**Part B** of **Fig. 1**).

The next reproduction cycle starts at generation *t^∗^* + 1 (**Part C** of **Fig. 1**), for a total of specified number of cycles *n_cycles_*, with final generation occurring before migration, i.e. *t _final_* = *n_cycles_t^∗^ −* 1. Visual representation of the experimental design of recombination rates and modes of reproduction is seen in **Fig. 2**

**Fig. 2.**
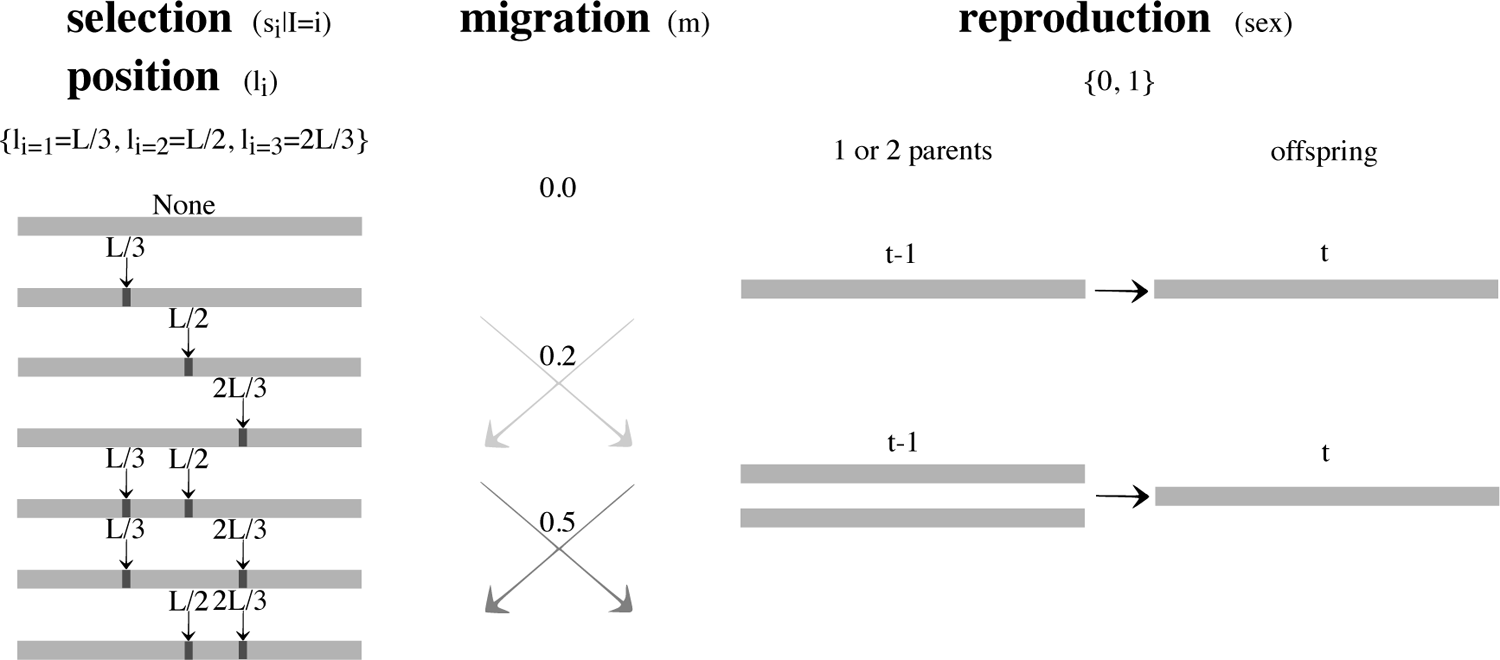
Possible parameter space of selection coefficients without the initial outlier scan on summarized observed data. Scheme of selection coefficients conditional on genome loci (*s_i_ I* = *i*), rates of migration (*m*), and possible modes of reproduction during migration generations (*sex*).

## 3 Model parametrization, the data, and the divergent selection simulator

In this section, we delineate the constraints on the data-generating process, define the parameters of the population genetic model, and finally build a simulator. The divergent selection simulator allows us to explore the divergent selection model motivated by the yeast populations built in the lab and it constitutes one of our main contributions.

### 3.1 Model parametrization and the data

To capture differences in population ***X*** relative to population ***Y***, we economically use signed selection coefficients. We arbitrarily fix the reference allele *a_i_* at locus *i* with selection coefficient *s_i_* (*i ∈* {1, 2, . . ., ∥***L_s_***∥}) for the other allele at that locus in population ***X***. A negative *s_i_* means that for allele *a_i_* at locus *i*, an individual in population ***X*** has a lower fitness in comparison to an individual in population ***Y*** . Unlike in population ***X***, the fitness of the reference allele *a_i_* under selection in population ***X*** is always (1 + 0) for a carrier in population ***Y***, regardless of whether the carrier in population ***Y*** has the *a_i_* copy or not.

SNPs in yeast occur for approximately one in every 165bp. We assume equally spaced SNPs in the genome and rescale the genome-wide recombination rate proportionally. We further assume that there is at most one crossover event between consecutive SNPs. We let *SNP_spacing_* be the spacings between *L* SNPs, which allows us to denote the total genome length by *L × SNP_spacing_*. The recombination rate is a function of *L* and *SNP_spacing_* is a constant: *r* = *f* (*L*), *SNP_spacing_* = *C*.

In the experimental laboratory setup, the migration rate and the length of migration-selection cycles *t^∗^* are controlled, and the recombination rate can be estimated as described earlier in the section 1, and in subsection 2.1. The migration rates are fixed at {0, 0.2, 0.5}. For the recombination rate *r*, we use values from the literature and assume them fixed and known. A typical computational procedure mimicking sequenced yeast genome informed by our laboratory procedures is as follows. We picked to simulate *L* = 1, 500 SNPs because it is computationally scalable. This number translates into about one-third on mean of SNPs of a chromosome that the two yeast strains in our biological model system from subsection 2.1 differ by over 70, 000 SNPs (Hether 2016). The mean recombination rate of *Saccharomyces cerevisiae* has been estimated as 3.5 *×* 10*^−^*^6^ Morgans/bp in the literature (Ruderfer et al. 2006), with inferred genome-wide recombination profiles from sequenced isolates from an advanced intercross line (AIL) to be as high as 3.0 *×* 10*^−^*^5^ Morgans/bp for a two-way cross at genome hotspots (Illingworth et al. 2013). Here, for our laboratory procedures mimicking the sequenced yeast, we fixed the genome recombination rate of *r* = 2.0 *×* 10*^−^*^5^ Morgans/bp. One way to think about this value is as a best statistical estimate for recombination hotspots in yeast. For the other scenarios with a smaller number of SNPs for computational time efficiency, we scaled the recombination rate to *r* = 3.0 *×* 10*^−^*^4^, proportionally to the expected number of recombination events.

Fixing unknown parameter values to their point estimates instead of jointly estimating them is not ideal. Recombination rate is known to be a particularly problematic parameter in population genetic models with selection and recombination. This is due to the fact that the two evolutionary processes might generate statistically similar signals in genetics. In models the recombination rate *r* was varied in our simulations the signal from divergent selection was confounded to the degree that there were no useful statistical inferences to distinguish true loci under selection. However, our work is a first serious effort to model divergent selection and to explore the statistical properties of estimates of a parameter of interest (strength of divergent selection) in a large evolutionary model. Free model parameters of not direct interest such as recombination rate cause statistical identifiability issues that jeopardize the statistical inference. These issues have not been solved in population genetics (except in small models) and their discussion is beyond the scope of this paper.

Statistical identifiability of model parameters kicks in when there is a large number of interacting parameters in a population genetics model that are not fixed, but vary.

**Table 1** describes the full parametrization with assumed values for fixed and known parameters and prior distributions for unknown and to be estimated parameters. We tried to choose reasonable support for the prior distributions based on yeast literature ((Bast et al. 2019; Hamil- ton et al. 2005; Boitard et al. 2016; Tsai et al. 2010; Illingworth et al. 2013; Hermann et al. 2019)). To be explicit in probability functions of the model representation, in addition to standard conditionality notation separating the observables and parameters, we denote the fixed and known parameters by ***K*** = (*r*, *t^∗^*, *n_cycles_*, *N_e_*, *L*, *SNP_spacing_*) where *r* is the recombination rate per genome per generation, *t^∗^* is the number of generations between reproduction cycles, *n_cyclec_* is the number of reproduction cycles, *N_e_* is the effective population size, and *L* is the number of SNPs, *SNP_spacing_* is the spacing on the genome between each SNP. We write the joint probability mass function generating the data as

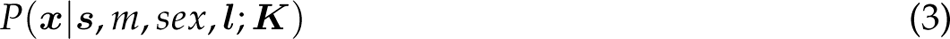

where, ***x*** is an *N_e_* by *L* matrix of zeros and ones for the SNP data for each population, ***s*** = (*s*_1_, *s*_2_, …, *s_I_****_Ls_****_I_*) is the vector of signed selection coefficients at each locus, *m* is the symmetric migration rate per generation between the two populations, *sex* is the mode of reproduction, ***l*** = (*l*_1_, *l*_2_, …, *l_I_****_Ls_****_I_*) is the vector of indicators to denote loci under selection.

**Table 1.** Properties of parameters in the model.

| Parameter | Status | Value | Prior | Motive |
| --- | --- | --- | --- | --- |
| $SNP_{spacing}$ | fixed and known | $\frac{\text{genome length}}{\text{number of SNPs strains differ by}} = 165$ | — | genome length of about 12 Mpb and strains differ by about 73,000 SNPs (Hether 2016) |
| $\mu_{nr}$ | fixed and known | 4.95 | — | expected value of number of crossovers = $\mathbb{E}(\text{Bin}(L \times SNP_{spacing} - 1, r))$ |
| $r$ | fixed and known | $2.0 \times 10^{-5}$ Morgans/bp, and $3.0 \times 10^{-4}$ Morgans/bp for $L = 1,500$ and $L = 100$ respectively | — | already done in literature (Tsai et al. 2010; Illingworth et al. 2013; Hermann et al. 2019) |
| $l_i$ | fixed, but unknown for method 1; not fixed and unknown for method 2, where potentially identified within outlier scan, specified by $set_{l_i}$ | — | $set_{l_i} = \{l_{i=1} = L/3, l_{i=2} = L/2, l_{i=3} = 2L/3\}$ for method 1; $set_{l_i}$ identified by initial outlier scan of 5% for method 2 | fixed loci as discrete value parameters at which selection acts to reduce the parameter space to $\{l_{i=1} = L/3, l_{i=2} = L/2, l_{i=3} = 2L/3\}$ and setting the values for rest of loci to zero for method 2; less selection parameter values $s_i \vert I = i$ to estimate in ABC after initial outlier scan, and setting the values for loci not identified by outlier scan to zero for method 2 |
| $s_i \vert I = i$ | unknown and estimated | — | $\text{Unif}[-0.25, 0.25]$ for $t^* = 5$, and $\text{Unif}[-0.025, 0.025]$ for $t^* = 50$ respectively | considered a large range of parameter space for $s_i \vert I = i$ in yeast (Gallet et al. 2012), and 10-fold smaller for 10-fold longer selection generation cycles for comparison |
| $n_r$ | random and unknown | — | — | number of recombination events sampled from $\text{Bin}(L \times SNP_{spacing} - 1, r)$ |
| $L_s$ | fixed but unknown | — | $\text{Discrete Unif}[set_{l_i}]$ | vector $L_s$ of length 0, 1, or 2 of loci under selection to reduce the parameter space |
| $L_r$ | random and unknown | — | — | vector $L_r$ of length $n_r$ of loci of recombination events sampled from $\text{Discrete Unif}[n_r, L \times SNP_{spacing} - 1]$ |
| $m$ | fixed but unknown | — | $\text{Discrete Unif}[n_r]$, $set_m = \{0.0, 0.2, 0.5\}$ | methods of estimation of $m$ already developed (Hamilton et al. 2005) but considering some variability ($set$) |
| $N_e$ | fixed and known | 10,000 | — | already done in literature (Tallmon et al. 2008; Boitard et al. 2016) |
| $L$ | fixed and known | 1,500 or 100 | — | 1,500 SNPs is about one-third on average of SNPs of a chromosome for a yeast cross (Hether 2016); 100 SNPs is for faster computational time |
| $t^*$ | fixed and known | 5 or 50 | — | short enough selection generation cycles of $t^* = 5$ to consider large range of parameter space for $s_i$; longer selection generation cycles of $t^* = 50$ for comparison |
| $n_{cycles}$ | fixed and known | 2 or 4 | — | two different population mixing frequencies for comparison on the signature of selection |
| $sex$ | fixed but unknown | — | $\text{Discrete Unif}[set_{sex}]$, $set_{sex} = \{0, 1\}$ | yeast system can reproduce sexually or asexually (Bast et al. 2019) |

### 3.2 Simulator for the data generating process

Below we describe how the data are generated with the simulator. We input effective population size *N_e_*, number of SNPs per carrier to be simulated *L*, loci corresponding to SNPs under selection ***L_s_***, alleles under selection *a*, recombination rate *r*, selection coefficients at loci under selection *s*, migration rate between populations *m*, number of generations between selection-migration cycles *t^∗^*, type of reproduction during migration generations *sex*, spacing of SNPs on the genome, assumed equal spacing *SNP_spacing_*, and number total number of generations to simulate *t _final_*.

The model parameters that are estimated from the output simulator are the loci under selection and corresponding selection coefficients, although we allow the migration rate between populations *m* and the type of reproduction during migration generations *sex* to vary between simulations but we do not estimate them. The estimated model parameters are sampled from the joint prior distribution, i.e. ***θ_i_^∗^*** *∼ P*(***s***, *m*, *sex*, ***l***; ***K***).

Our simulated data of bi-allelic SNPs are *N_e_ × L* matrices per population represented by *x*. The SNPs of each carrier in the population correspond to the matrix row. For each of *N_e_* carries per population an allele *a* is sampled from discrete uniform distribution bound on [0, 1] and replicated *L* times, corresponding to the probability of 0.5 of carriers from founding *F*_0_ population ***X*** and of 0.5 from founding *F*_0_ population ***Y***, to build *F*_1_ populations ***X*** and ***Y*** respectively.

Offsprings are then generated from the recombination of two parental genomes per offspring. For 1 to *N_e_* per population, the first parent is sampled with equal probability from ***X_F_*_1_**, and the second parent is sampled with equal probability from ***Y_F_*_1_**, both from a discrete uniform distribution, i.e. *p*_1_ *∼* Discrete Unif[1, *N_e_*] from ***X_F_*_1_** and *p*_2_ *∼* Discrete Unif[1, *N_e_*] from ***Y_F_*_1_** . The number of recombination events *n_r_* between *p*_1_ and *p*_2_ are sampled from *Bin*(*L × SNP_spacing_ −* 1, *r*), with *r* corresponding to genomic recombination rate and *SNP_spacing_*corresponding to SNP spacing on the genome. Loci of recombination events, ***L_r_***, are sampled *n_r_* times from ***L_r_*** *∼* Discrete Unif[*n_r_*, *L × SNP_spacing_ −* 1]. If a random variable *rv* sampled from Uniform distribution on [0, 1] is less than 0.5, i.e. if *rv ∼* Unif[0, 1] *<* 0.5, *p*_1_ is reassigned to *p*_2_ and *p*_2_ is reassigned to *p*_1_. The recombination of *p*_1_ and *p*_2_ then starts from the genome of *p*_1_ and alternates between two parents to form an offspring. This concludes founding *F*_2_ at time *t* = 0.

Then, the selection and asexual reproduction occur for *t^∗^ −* 1 generations. Absolute fitness of parents 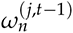 for each population *j* in ***X***, ***Y*** for each carrier *n* is calculated. For loci under selection specified by an input vector ***L_s_***, absolute fitness of parent at *t −* 1 is 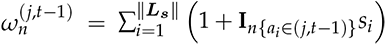, where *s_i_* correspond to non-zero selection coefficients specified by vector 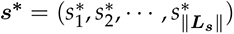, and alleles under selection specified by vector 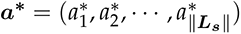.

Probability of carrier in the population *p_n_* having an offspring is the normalized absolute fitness 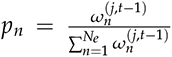. The *N_e_* offspring are 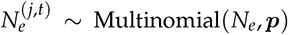, with normalized absolute fitness probabilities ***p*** = (*p*_1_, *p*_2_, . . ., *p_Ne_*).

At generation, *t^∗^* migration between populations takes place, specified by migration rate *m*, with the corresponding type of reproduction specified by *sex ∈* {0, 1}.

The selection-migration cycles as described in **Part A** and **Part B** respectively are repeated for a total specified number of cycles *n_cycles_*, with ending on the final generation before migration, i.e. *t _final_* = *n_cycles_t^∗^ −* 1. The output of the data of the *x* matrices of dimension *N_e_ × L* per population is then ready for model inference by mapping the data to summary statistics and the ABC.

## 4 Inference about model parameters via Approximate Bayesian computation (ABC)

The probability distribution given in expression 3 is not available in closed form and the joint likelihood of the data cannot be evaluated given the parameters. There are no exact methods to perform statistical inference about the unknown model parameters in this case. As a practicable solution to the problem of performing inference about the model parameters, we employ ABC to sample the posterior distribution of parameters (Li & Jakobsson 2012; Quinto-Cortés et al. 2018; Smith & Flaxman 2020; Beaumont et al. 2002; Estoup et al. 2004; Fagundes et al. 2007; Cornuet et al. 2014; Sousa et al. 2012). ABC bypasses the explicit evaluation of the joint likelihood thereby making simulation-based inference feasible when model likelihoods cannot be evaluated. Statistical inference in ABC is characterized by two main approximations. The first approximation is due to substituting the exact likelihood of the data with a kernel-based numerical approximation. The second approximation is due to substituting the likelihood of the data with the likelihood of the summary statistics. Whether and by how much these approximations affect the quality of the inference depends on the size of the model generating the data and the computational budget available to increase accuracy of the approximation. For the first approximation, good practices have been established. Assessing the quality of the second approximation, however, is particularly challenging in a class of models where there are no known sufficient statistics for unknown parameters. Our divergent selection model falls into this class. In this section, we investigate the usefulness of some population differentiation statistics to perform inference about the parameters of our divergent selection model.

Based on expression 3 for the probability model generating the data, we denote the joint posterior distribution of parameters given the data ***x*** and fixed and known parameters as

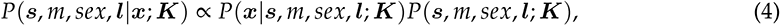

where *P*(***x****|****s***, *m*, *sex*, ***l***; ***K***) is the joint likelihood of the data and *P*(***s***, *m*, *sex*, ***l***; ***K***) is the joint prior distribution of unknown parameters. Incorporating the two approximations described in the previous paragraph, we write the likelihood as

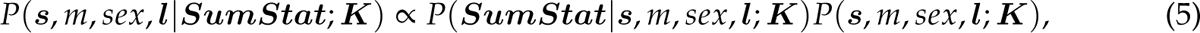

where ***SumStat*** is summary statistics. In the next two subsections, we evaluate some useful summary statistics in the context of our model.

### 4.1 Summary Statistics and Outlier Scan on Summarized Observed Data

The effect of divergent selection on genomes between samples of two populations can be quantified by well-known statistics that measure of genetic differentiation. An example is Wright’s fixation index (Wright 1949), *F_ST_*. For bi-allelic loci *F_ST_* defined by:

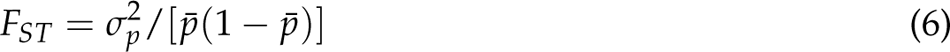

which measures allele frequency differentiation among the sampled populations (Holsinger & Weir 2009). Here, *σ_p_*^2^ is the variance in allele frequency among sampled populations, and *p̄* is the mean allele frequency in sampled populations. A signed version of *F_ST_*, which we denote by *signF_ST_* obeys the following: **if** (*p_X_ − p_Y_*) *<* 0, *signF_ST_* = *−F_ST_*, **else** *signF_ST_* = *F_ST_*, such that *signF_ST_ ∈* [*−*1, 1]. This statistic captures the information about which sampled population is undergoing selection advantageous with respect to the other sampled population.

At the genomic scale, it is often computationally infeasible to approximate the likelihood based on *F_ST_* jointly for all loci. A practicable remedy, which we follow here, is first to determine a set of candidate loci under selection manifesting only outlier values of *F_ST_* and consider the likelihood based on these loci. We take the outlier cutoff to be *F_ST_* outside of the 95% of all *F_ST_* values in the data (Beaumont & Nichols 1996). The selection coefficients can take on values within a range defined in **Table 1**, it is not a single value, for which we can derive an estimated *F_ST_* cut-off value, therefore we do not consider just large *F_ST_* values. In the simulator, the summary statistics outliers correspond to specific SNPs. Only the specific loci of SNPs that were detected by the outlier test are considered as potentially under selection when simulating data sets for ABC. The rest of the loci (non-outliers) are fixed to as no selection, i.e. *s_i_* = 0.

Sample *F_ST_* is not a sufficient statistic for any parameter of a divergent selection model since large values of sample *F_ST_* are not only the result of divergent selection. Sample *F_ST_* from expression 6 measures genetic variation among sampled populations by assessment of variance between and within sampled populations by calculating allele frequency differences. The sample *F_ST_* values have been found to be correlated with the recombination rate (Laayouni et al. 2011). Recombination rate between loci and the number of generations of recombination influence LD decay with genetic distance. Under a neutral evolution model, genetic drift is the only driving force of changes in allele frequencies. Many generations will be required for a new variant to reach a high frequency, and the surrounding LD will decay due to recombination events (Kimura 1983; Otyama et al. 2019; Bomba et al. 2015).

In our paper we test an LD-based summary statistic – Cross Population Extended Haplotype Homozygosity (*XP − EHH*) (Sabeti et al. 2002; Weigand & Leese 2018) – along with the sample *F_ST_*. In order to derive sample *XP − EHH*, one must first calculate Extended Haplotype Homozygosity (*EHH*) summary statistics for each population.

The *EHH* between two loci is defined as the probability that the number of distinct haplotypes *G_v_* in a genomic region up to a distance *v* from the locus are equal to each other. For *N_e_* carriers per population with possible alleles of either 0 or 1 per locus, with each group *z*, *z* = 1, 2, . . ., *G_v_*, having *n_z_* haplotypes, *EHH* is:

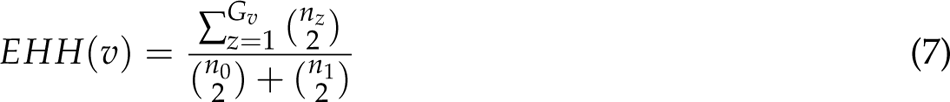

(Sabeti et al. 2007). At distance, *v* = 0 the *EHH*(*v* = 0) for each locus with respect to itself is always 1 and the *EHH*(*v*) values decay as the *v* increases, which is the decay of LD from each core haplotype (Sabeti et al. 2002; Voight et al. 2006). The *XP − EHH*, compares the integrated extended haplotype homozygosity, *EHH*, between two populations (Sabeti et al. 2002, 2007). Specifically, *XP − EHH* is the ratio of the *EHH* between populations ***X*** and ***Y*** integrated over the genome. If recombination rates in the model were allowed to vary widely across the genome between and within populations, the EHH statistic can be interpreted as a measure of selection only after suitable normalization (Sabeti et al. 2007; Wagh et al. 2012). Our model assumes a mean recombination rate for simulated *L* SNPs, thus normalization is not required in our model.

Calculating *EHH*(*v*) values for locus 1 to *L*, and corresponding distances *v* 0 to *L −* 1, it yields an *L* by *L* symmetric matrix for each of the two populations. The *XP − EHH* then for population ***X*** and ***Y*** combined for each locus is just a vector of length *L*, and is given by:

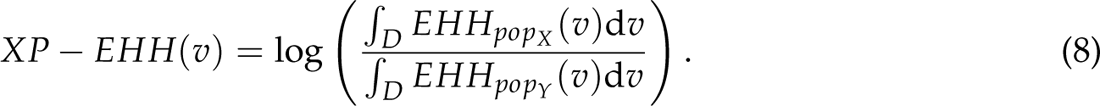

The integration domain *D* is a cut-off threshold below which the *EHH* values are set to 0.0, for which we pick *D* of 0.05 (Voight et al. 2006; Qanbari et al. 2011). The XP-EHH has the advantage of detecting selection on alleles that near fixation in one population but not both, population ***X*** and ***Y*** (Sabeti et al. 2007). This fits our model where the population ***X*** evolves under the positive or negative selection with respect to population ***Y***, with occasional gene flow between two populations during migration generations.

### 4.2 Assessment of Summary Statistics

Following the outlier scan on the summarized observed data set (*F_ST_*statistic) outside of the 95% of all *F_ST_* values as described in the previous subsection, only the outlier loci in the data sets simulations identified by the outlier scan are considered as potentially under selection, with selection coefficients at non-outlier loci fixed to 0, and only the summary statistics corresponding to loci identified by outlier scan are inputted to **Algorithm 1** and **Algorithm 2**. We assess the performance of summary statistics in terms of how well they capture the signal of observed selection coefficients by using the summary statistics in the ABC, then calculating the standard deviation of posterior distributions of selection coefficients, and mean square errors (MSEs), variance and squared-bias. We compare performance of the following summary statics: *signF_ST_*, *XP − EHH*, *p_X_ − p_Y_*, and *signF_ST_* with *XP − EHH*. We plot SNPs vs. MSE, observed selection coefficients vs. MSE, SNPs vs. squared-bias, observed selection coefficients vs. squared-bias, SNPs vs. variance, observed selection coefficients vs. variance, for the four summary statistics combinations, and for the mode of reproduction and migration rate in section 5.

In our model *p_X_* and *p_Y_* represent allele frequency per locus in sampled population ***X*** and ***Y*** respectively, such that *p_X_ − p_Y_*is the difference between the allele proportions per locus between the two populations, with the expected value of zero for the founding *F*_2_ populations, i.e. at generation *t* = 0. Additionally to testing the performance of summary statistics from the outlier scan on summarized observed data described above, we assessed the effect of genetic drift and found that the population size is large enough for genetic drift not to be an issue as the mean of 100, 000 data sets converge to the deterministic model, as verified by simulations. Due to the convergence of the mean of 100, 000 data sets and the expected value from the deterministic model, we determined most informative summary statistics performed of simulator output data (*t* = *t _final_*) on the deterministic model based on the MSEs. We derived a summary statistic called *signF_ST_*: **if** (*p_X_ − p_Y_*) *<* 0, *signF_ST_* = *−F_ST_*, **else** *signF_ST_* = *F_ST_*. Due to the strong fit of superimposed plots of simulations and deterministic values (Gillespie 2004), we evaluated *signF_ST_*, *F_ST_*, and (*p_X_ − p_Y_*) based on deterministic single locus model. We performed 1,000 ABC tests with a tolerance rate of 0.1%. For each of the ABC iterations, a single simulated data set with known parameters was randomly drawn from the 100, 000 data sets and assumed as observed data set. For each, the top 0.1% of data sets with the smallest euclidean distances between observed and simulated summary statistics were accepted. On average of the 1,000 ABC iterations, the lowest error was achieved with *signF_ST_* (*MSE* = 5.50 *×* 10*^−^*^8^). We compared ABC results to empirical results from the simulator from 100,000 simulations and achieved on average *MSE* = 2.00 *×* 10*^−^*^3^ with *signF_ST_* and *MSE* = 1.10 *×* 10*^−^*^2^ with (*p_X_ − p_Y_*). We also examined parameter space via plots of *s_L_*_/2_ vs. summary statistics and the relations are more often 1:1 with *signF_ST_* than with (*p_X_ − p_Y_*). In section 5 we address *XP − EHH* and *signF_ST_* and expand it to a much larger scope of parameter space.

### 4.3 Inference About Model Parameters

We estimate the free model parameters in *P*(***s***, *m*, *sex*, ***l****|****SumStats***; ***K***) given by expression 5, using an ABC-rejection algorithm (**Algorithm 1**) and the ABC with the linear regression adjustment (**Algorithm 2**). The term ***x*** represents, the data from our bi-allelic SNPs of individuals in two populations. The simulated *SNPs* are *N_e_ × L* matrices per population represented by ***x***, and ***x*** summarized by summary statistics *SumStat* by mapping ***SumStat*** = *S*(***x***). The *nsim* observations denoted by ***x*** generated from the model are independent and identically distributed, i.i.d. The ***x*** *∈ X*, where *X* is the space in which the data sits.

ABC makes two approximations, the first one is the mapping of ***x*** to ***SumStat***, and the second is accepting summary statistics ***SumStat*** within a tolerance rate from the observed ***SumStat_obs_***. The simulations are a mechanistic process that involves random sampling, which can be thought of as an influence of stochastic processes such as genetic drift. The ABC facilitates in model parameters estimation by accepting parameters corresponding to summarized data sets within a tolerance rate that is partly due to the stochastic effect. The ABC outputs a posterior distribution of accepted parameters *P*(***s***, *m*, *sex*, ***l****|****SumStat_m_***; ***K***), *m ∼* Discrete Unif[*set_m_*], where *set_m_ ∈* [0, 1]. In the ABC-rejection (**Algorithm 1**) we calculate the Euclidean distance *d_i_*, *i* = 1, 2, . . ., *nsim* technique for each of the simulated and summarized by the summary statistics data set (Greenacre 2017; Prangle et al. 2017) to scale summary statistics across the *nstat* dimensional space. The summary statistics are standardized by the median absolute deviation *SD*, *j*, *j* = 1, 2, . . ., *nstat*, such that each of the *nstat* summary statistics per dimension approximately equally contribute to the ABC analysis. Further details on the Euclidean distance are described in **Appendix A**.

#### Algorithm 1

ABC-rejection algorithm for summary statistics calculated from data sets simulated from the **Simulator** and their corresponding parameters from the prior distribution. Input: proportion of simulations to accept, number of simulated data sets, number of summary statistics per simulation

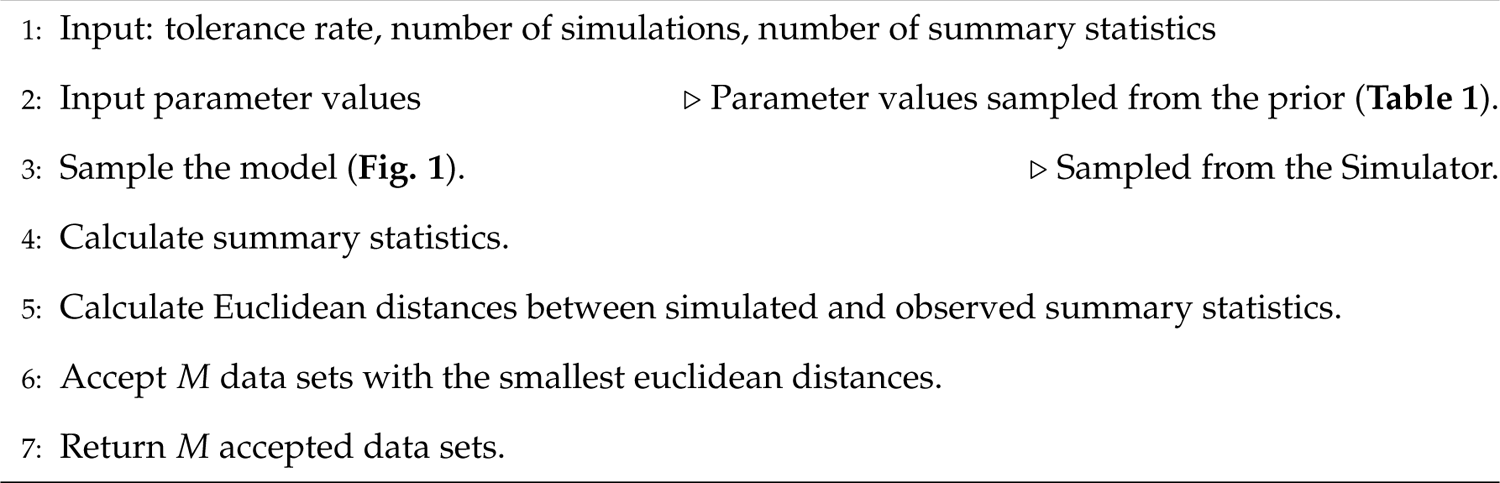

Following the rejection algorithm, we applied the linear regression correction to compare ABC performance with and without linear regression correction. The input of the ABC-linear regression is the output of the ABC-rejection. In ABC-rejection we assign weights *w_m_* to accepted data sets from ABC-rejection (Park et al. 2016). The weight *w_m_* for each of accepted pairs from **Algorithm 1** output are calculated using kernel κ*_dM_* (*d_m_*) (Nott et al. 2014), where *d_m_* is the *m^th^* smallest euclidean distance between standardized accepted summary statistics (***SumStat^∗^ scaled***) and standardized observed summary statistics (***SumStat_obsscaled_***), and *d_M_ >* 0 is the bandwidth parameter (Blum 2017), in this case the the largest of *M^th^* euclidean distance, order *(d*_1_, *d*_2_, *· · ·*, *d_nsim_)* [*M*]. The purpose of the kernel weights calculations is to apply in the calculation of weighted least squares regression coefficients ***β***^^^*_W LS_* for the linear regression correction analysis in **Algorithm 2**, where the accepted data set with smallest euclidean distance *d_i_*from **Algorithm 1** output is adjusted the least and the accepted data set with largest *d_i_*out of the *M* accepted is adjusted the most. The beta estimates vector is an approximate draw from the posterior. In our ABC linear regression correction, the user can choose to calculate the kernel weights either based on Gaussian (Park et al. 2016; Csilléry et al. 2010), or Epanechnikov (Epanechnikov 1969) kernel and we later show in the results that the two kernels are similarly as effective. Further details on kernels, kernel weights, weighted least squares coefficients, and adjustment of accepted data sets parameter estimates are described in **in Appendix A**.

The ABC-linear regression correction steps are shown in **Algorithm 2**.

#### Algorithm 2

ABC-linear regression correction algorithm performed on the output of the ABC-rejection. Input: standardized observed summary statistics, *M* accepted standardized summary statistics with corresponding *M* accepted parameters from ABC-rejection.

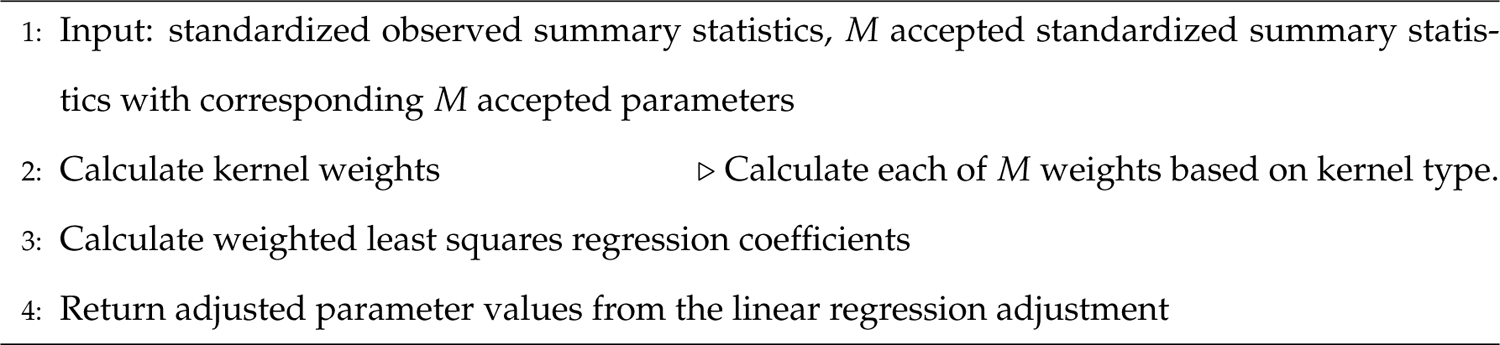

## 5 Results

### 5.1 Overview

Below we describe two methods for estimating strength of selection under four migration (*m*) and mode of reproduction (*sex*) combinations, and 0,1, or 2 loci under selection combinations.

In the first method we describe, *t^∗^* and *t _final_* are fixed 5 and 19 respectively, and the selection coefficients conditional on locus span on *s_i_|I* = *i ∼* Unif[-0.25,0.25] (see subsection 3.2). The parameter space for loci under selection is at genomic locus corresponding to SNP number that can only take on values on {*L*/3, *L*/2, 2*L*/3}. An advantage for this method is no need for the initial summary statistic outlier scan (as described in subsection 4.1) which would require simulation of *n* = 100, 000 data sets per observed data set, with parameter space of potential loci under selection reduced to those identified by the outlier scan. In subsection 5.2 and sub-subsection 5.2.1 we describe and perform an ABC analysis for each of total of *n_ABC_* = 10, 000 observed data sets, where we re-use simulated same *nsim* = 100, 000 data sets for each analysis. We plot observed *signF_ST_* summary statistics for all 10, 000 observed data sets and visually represent how the parameter space for the strength of selection differs depending on migration and mode of reproduction combinations (**Fig. 3**), and depending on number of loci under selection (**Fig. 4**). To reduce the complexity of variable parameters, we visually verify of this, simplified method case that ABC-linear regression correction estimation outperforms ABC-rejection estimation, and that there is no difference in performance of ABC-linear regression with Gaussian kernel versus with Epanechnikov kernel (**Appendix C**).

**Fig. 3.**
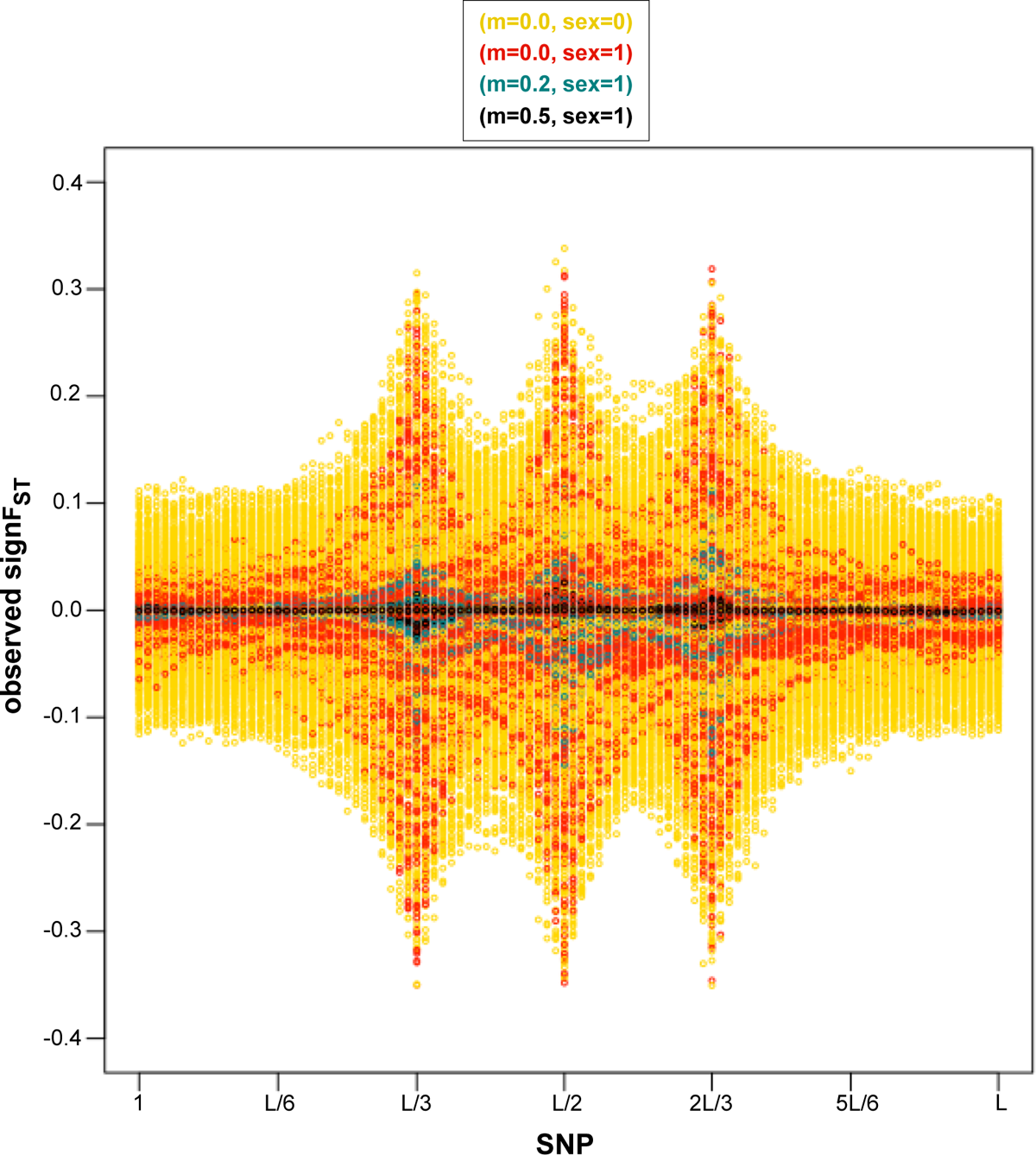
SNP loci vs. *signF_ST_* summary statistics for each of 10, 000 simulator output data sets assumed as observed under four of migration (*m*) and mode of reproduction (*sex*) combinations selected randomly with equal probability: *sex* = 0 and *m* = 0.0 in yellow, *sex* = 1 and *m* = 0.0 in red, *sex* = 1 and *m* = 0.2 in turquoise, *sex* = 1 and *m* = 0.5 in black, and selected randomly with equal probability scenarios of loci under selection as seen in **Fig. 2**. The visible pattern in an increase in summary statistics values away from 0 at selected loci with a decrease in migration rate, and increase in genetic hitchhiking effect with an asexual mode of reproduction (*sex* = 0) given same *m* of *m* = 0.0.

**Fig. 4.**
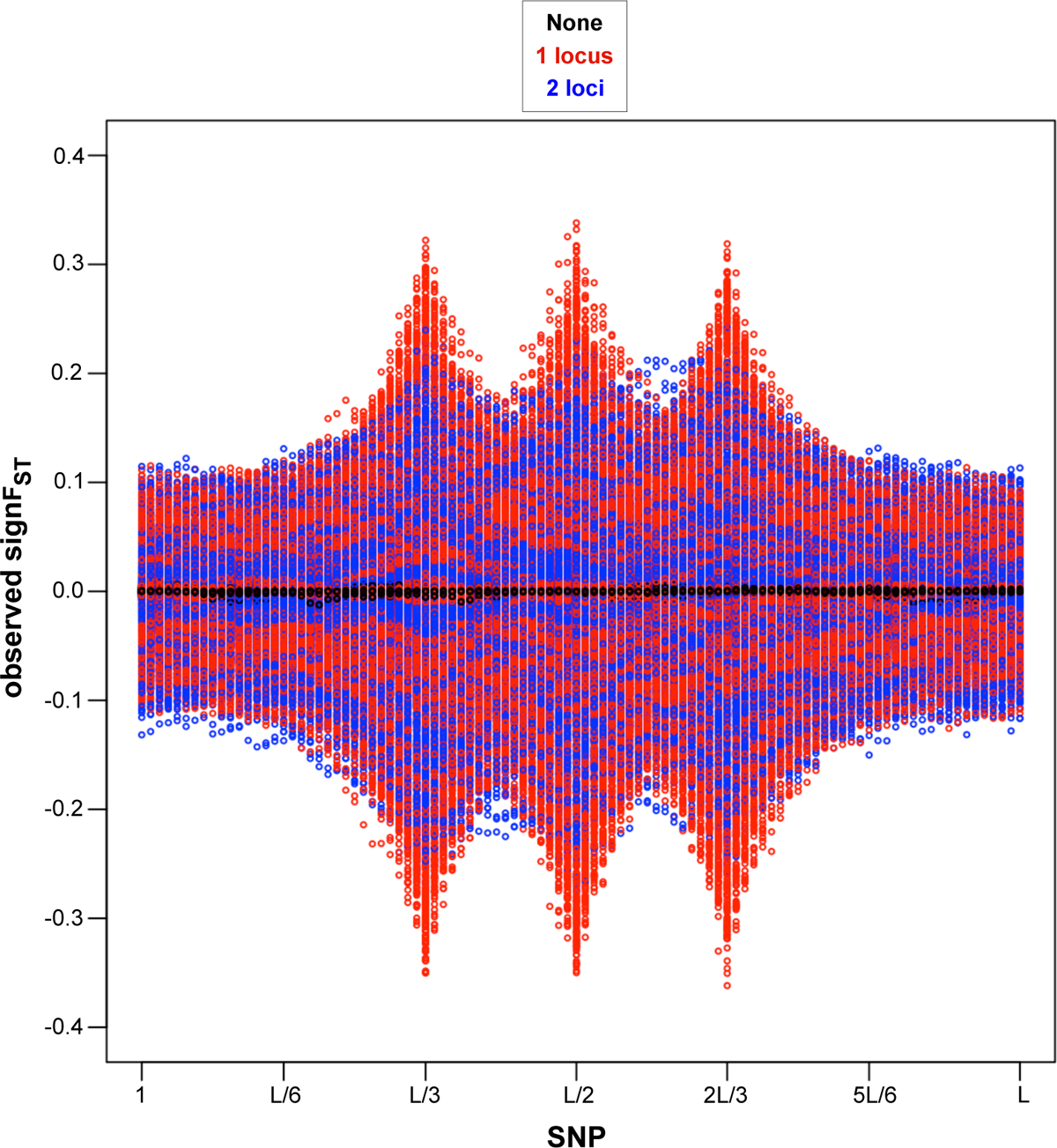
SNP loci vs. *signF_ST_* summary statistics for each of 10, 000 simulator output data sets assumed as observed under four of migration (*m*) and mode of reproduction (*sex*) combinations selected randomly with equal probability scenarios of loci under selection: none in black, one locus (*L*/3, or *L*/2, or 2*L*/3) in red, two loci in blue (*L*/3 with *L*/2, or *L*/3 with 2*L*/3, or *L*/2 with 2*L*/3), as seen in **Fig. 2**. Visible pattern in summary statistics values closest to 0 for no loci under selection, largest magnitude in summary statistics values away from 0 at single locus under selection, and in-between the magnitude of summary statistics values away from 0 and increase in genetic hitchhiking effect for two loci under selection.

In the second method, we performed four scenarios total of *t^∗^*, *t _final_*, number of SNPs *L*, recombination rate *r*, and parameter space of selection conditional on locus *s_i_|I* = *i* combinations (see **Table 2**), unlike where in first method we performed only one combination. Additionally, we build on the technique from the first method by allowing the loci under selection of the observed data to span anywhere between locus corresponding to SNP 1 to *L*, then performing outlier scan as as described in subsection 4.1 which requires simulation of *n* = 100, 000 data sets per observed data set. Based on the results from the first method about ABC-rejection versus ABC-linear regression, and corresponding ABC-linear regression two kernels, we performed analysis only with ABC-linear regression (after the ABC-rejection but without comparing to ABC-rejection) and compare ABC performance with four summary statistics combinations instead of one like in the first method, and found that *signF_ST_* summary statistics in ABC analysis is has low squared-bias and variance between observed and median accepted posterior selection parameters relative to the other three summary statistics combinations (**Fig. 9**).

**Table 2.** Scenarios under which the initial parameters varied for the model simulations and ABC, with parameter values in blue that are unique to particular scenario, and those in black that are shared with at least one other scenario.

|  | Scenario 1 | Scenario 2 | Scenario 3 | Scenario 4 (sequenced yeast data) |
| --- | --- | --- | --- | --- |
| **Varying Parameters** | $L = 100, r = 3.0 \times 10^{-4}, t^* = 5, t_{final} = 19, s_i\vert I = i \sim \text{Unif}[-0.25, 0.25]$ | $L = 100, r = 3.0 \times 10^{-4}, t^* = 50, t_{final} = 199, s_i\vert I = i \sim \text{Unif}[-0.025, 0.025]$ | $L = 100, r = 3.0 \times 10^{-4}, t^* = 50, t_{final} = 99, s_i\vert I = i \sim \text{Unif}[-0.025, 0.025]$ | $L = 1,500, r = 2.0 \times 10^{-5}, t^* = 50, t_{final} = 199, s_i\vert I = i \sim \text{Unif}[-0.025, 0.025]$ |
| **Total number of ABC iterations ($n_{ABC}$)** | 100 | 100 | 100 | 10 |
| **Total number of simulated data sets for all ABC iterations ($n_{ABC} \times nsim$)** | 10 million | 10 million | 10 million | 1 million |

### 5.2 Observed Data, Model Simulations, and ABC for Fixed Potential loci under Selection

In this first method, we assessed how the signal of selection from summary statistics changes when the mode of reproduction, strength of migration, as well as number of loci under selections change. As Wright’s fixation index (Wright 1949) measures the allele frequency differentiation among sampled populations (Holsinger & Weir 2009) of sequenced data, and demographic history of *Saccharomyces cerevisiae* has been reported to play a role in gene expansion and contraction based on phylogeny reconstruction (Duan et al. 2018), here we investigate this demographic history and its effect on signature of selection.

For this, we assessed signal strength equivalent to scenario 1 for the second method from **Table 2** but for 10, 000 ABC iterations (*n_ABC_* = 10, 000) and without the initial outlier scan on summarized observed data. This method has a computational time advantage as instead of simulating unique 100, 000 (*nsim* = 100, 000) data sets per one observed data, the simulated data sets are reused. We present the results with *signF_ST_* summary statistics based on ABC-linear regression with Gaussian kernel (see **Fig. 5**, and **Fig. 6**), but we also compare performance of Gaussian versus Epanechnikov kernels, Gaussian kernel versus rejection, and Epanechnikov kernel versus rejection (see **Appendix C**).

**Fig. 5.**
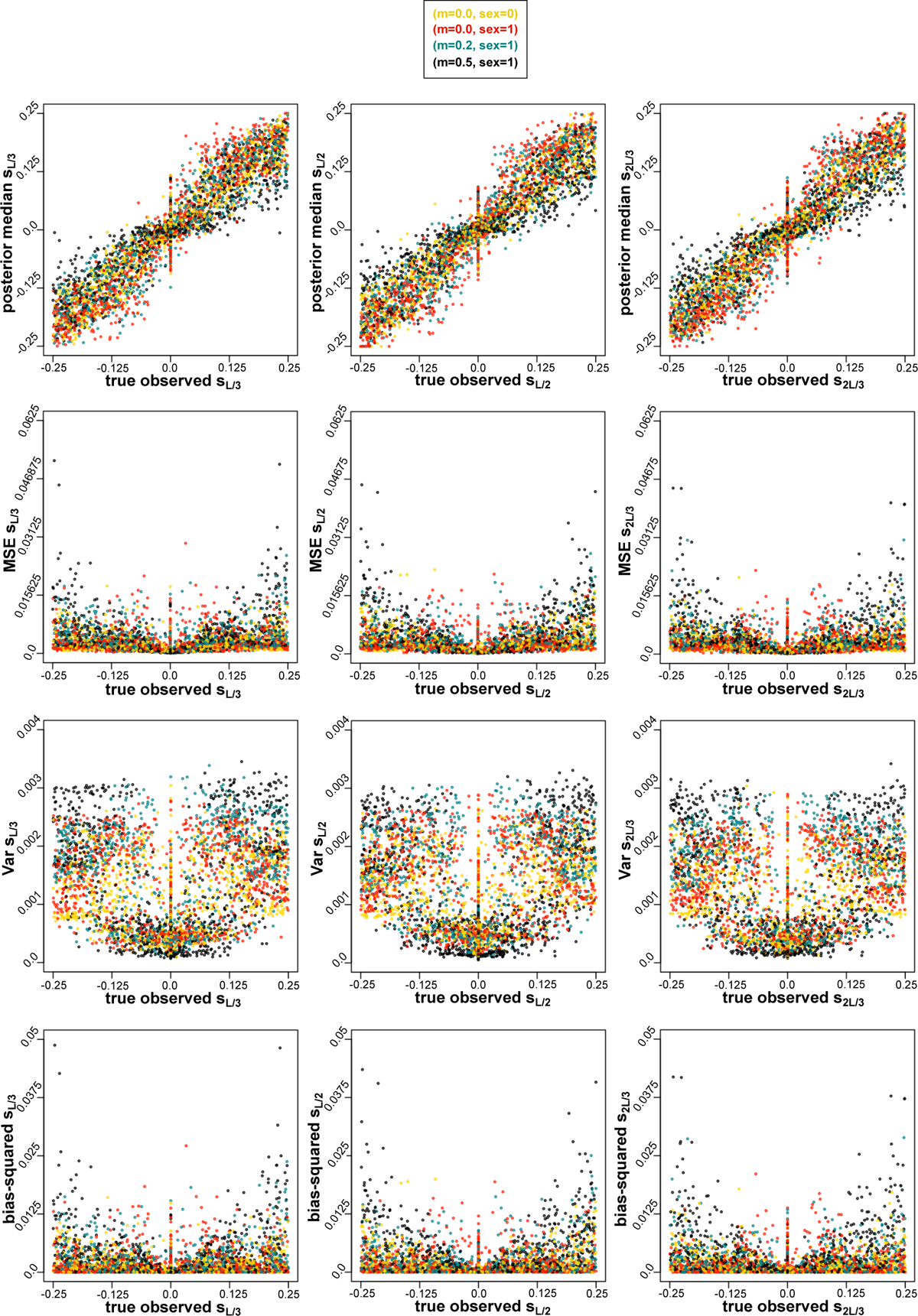
True parameter value under which the observed parameter is generated (x-axis) vs. median, MSE, variance, bias-squared for (*L*/3)^th^, (*L*/2)^th^ and (2*L*/3)^th^ SNP respectively from 10,000 ABC tests from *signF_ST_* summary statistics from **Algorithm 2** with Gaussian k3e1rnel. Colors based on four migration (*m*) and mode of reproduction (*sex*) combinations of observed data sets: *sex* = 0 and *m* = 0.0 in yellow, *sex* = 1 and *m* = 0.0 in red, *sex* = 1 and *m* = 0.2 in turquoise, *sex* = 1 and *m* = 0.5 in black. A positive relationship between the magnitude of selection for strongest migration rate (*m* = 0.5) and variance as well bias (squared).

**Fig. 6.**
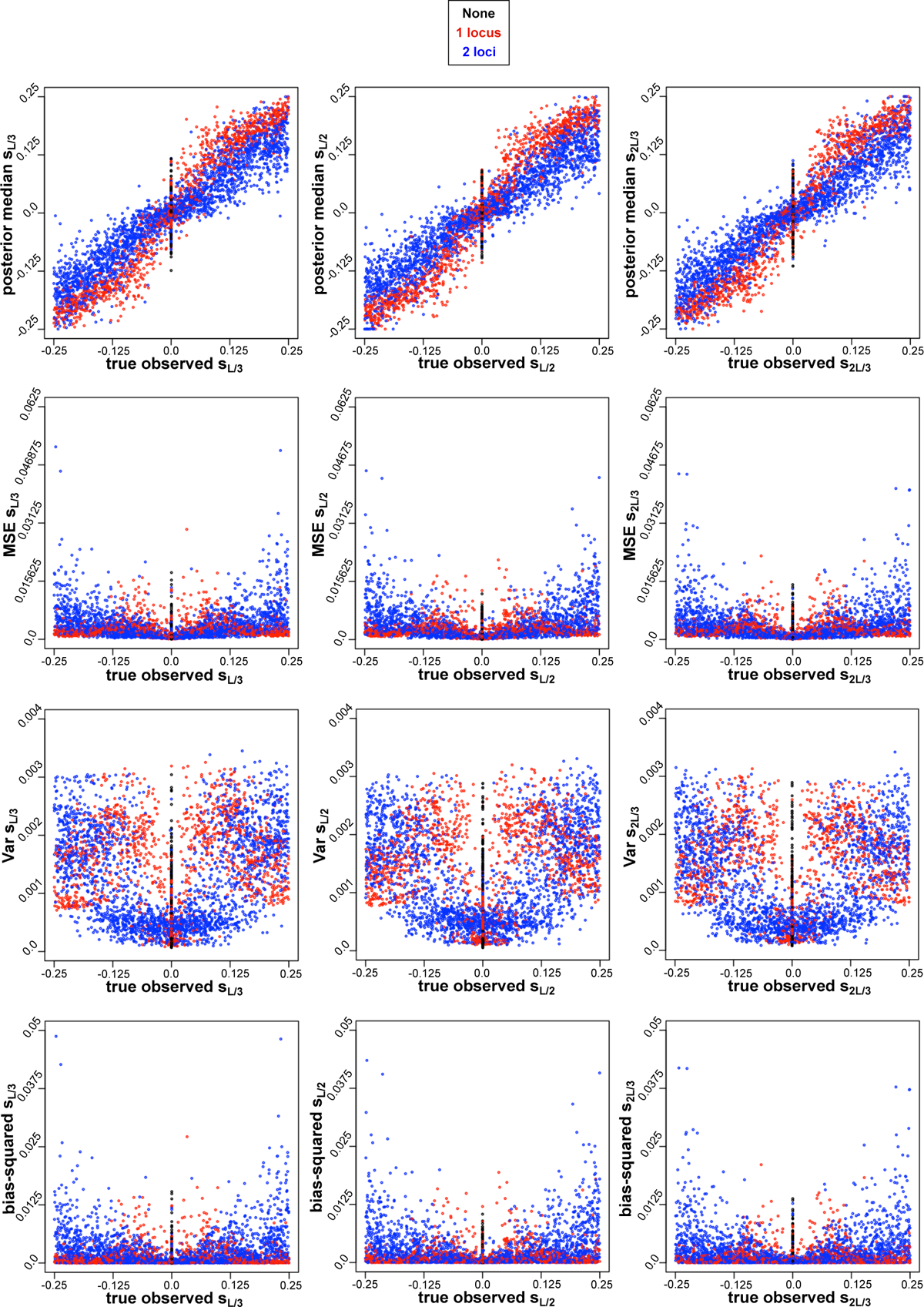
True parameter value under which the observed parameter is generated (x-axis) vs. median, MSE, variance, bias-squared for (*L*/3)^th^, (*L*/2)^th^ and (2*L*/3)^th^ SNP respectively from 10,000 ABC tests from *signF_ST_* summary statistics from **Algorithm 2** with Gaussian kernel. Colors based on three combinations of number of loci under selection of observed data sets: none in 3b2lack, one locus in red, and two loci in navy blue. Positive relationship between the magnitude of selection for two loci and variance as well bias (squared). Change from positive to negative relationship between the magnitude of selection for one locus and variance as well bias (squared) as the magnitude of selection increases.

The details of the experimental design of parameter space are described below.

#### 5.2.1 Experimental Design of Parameter Space

The experimental design of the parameter space which was the same as for the scenario 1 in **Table 2** is as follows: for *L* = 100 SNPs, with recombination rate of 3.0 *×* 10*^−^*^4^ Morgans/bp. The expected number of crossovers was *µ_nr_* = 4.95 and average spacing of polymorphic sites on the genome every *SNP_spacing_* = 165 sites as those we used in scenario 4 (**Table 1**, and **Table 3** in **Appendix A**), resembling a biological yeast system where YPS128 and DBVPG1106 yeast strains of 12Mb genome length differ by over 70,000SNPs or 1 SNP per *∼* 165 bp (Hether 2016), as described in subsection 2.1. The migration rate was randomly chosen to be 0.0, 0.2, or 0.5, sexual or asexual reproduction during migration generation. The migration generation cycles (*n_cycles_* = 4) took place every fifth generation (*t^∗^* = 5) for total of *t _final_* = *n_cycles_t^∗^ −* 1 = 19 generations. Both, the observed and simulated data can have either 0, 1, or 2 loci under selection with prior parameter space of selection coefficients conditional on locus *s_i_|I* = *i ∼* Unif[*−*0.25, 0.25].

**Table 3.** Description of conversion between genomic recombination rate, and recombination rate with respect to SNPs.

|  |
| --- |
| One crossover of actual genome that occurs between 2 consecutive SNPs actually would be one crossover between simulated SNPs only. Two crossovers between consecutive SNPs on the genome would look like no crossover between simulated SNPs, which we do not want to consider because it is very rare. Below is a note of how the recombination conversion is performed:<br>For yeast cross from the lab as example, genome length=12.066Mb and strains differ by 73,015 SNPs<br>$SNP_{spacing} = 12.066\text{Mb} / 73,015 \approx 165$<br>The recombination rate conversion inside the function:<br>$avg_{numrec} = (L \times SNP_{spacing} - 1) \times r$<br>(This is the average number of recombination events of genome chunk on which the SNPs are spaced)<br>Example: For $L = 100$, and $r = 3.0 \times 10^{-4}$<br>$avg_{numrec} = 4.95$<br>Convert $r$:<br>$r_{SNP} = avg_{numrec} / (L - 1) = 4.95 / 99 = 0.05$<br>(This is the recombination rate of SNPs alone)<br>$avg_{numrecSNP} = (L - 1) \times r_{SNP} = 99 \times 0.05 = 4.95 = avg_{numrec}$<br>(This is the average number of recombination events of SNPs alone)<br><u>Parameters per SNP equivalent to line 13 and line 15 of Algorithm 3:</u><br>Sample number of recombination events independently of each other (line 13):<br>$n_r \sim \text{Bin}(L - 1, r_{SNP})$<br>Sample loci of recombination events (line 15):<br>$\mathbf{L}_{r_{SNP}} \sim \text{Discrete Unif}[n_r, L - 1]$<br><u>What is written in line 13 and line 15 of Algorithm 3:</u><br>Sample number of recombination events independently of each other (line 13):<br>$n_r \sim \text{Bin}(L \times SNP_{spacing} - 1, r)$<br>Sample loci of recombination events (line 15):<br>$\mathbf{L}_r \sim \text{Discrete Unif}[n_r, L \times SNP_{spacing} - 1]$<br>The parents reproduce sexually and parental genome recombination takes place. We denote probability of genome recombination with $r$ (Morgans/bp units). We sample number of recombination events $n_r$ from Binomial distribution with a sequence of independent $L \times SNP_{spacing} - 1$ experiments with probability $r$. For each of the $n_r$ recombination events, we sample genome loci of recombination events $\mathbf{L}_r$, from Discrete Uniform distribution from locus 1 to locus $L \times SNP_{spacing} - 1$, but we assume no more than one recombination event between two consecutive SNPs. The cross of two populations at $t = 0$ (end **Part II** and start **Part A** of **Algorithm 3**) serves as an ancestral pool for $\mathbf{X}$ and $\mathbf{Y}$ populations. |

**Table 4.**
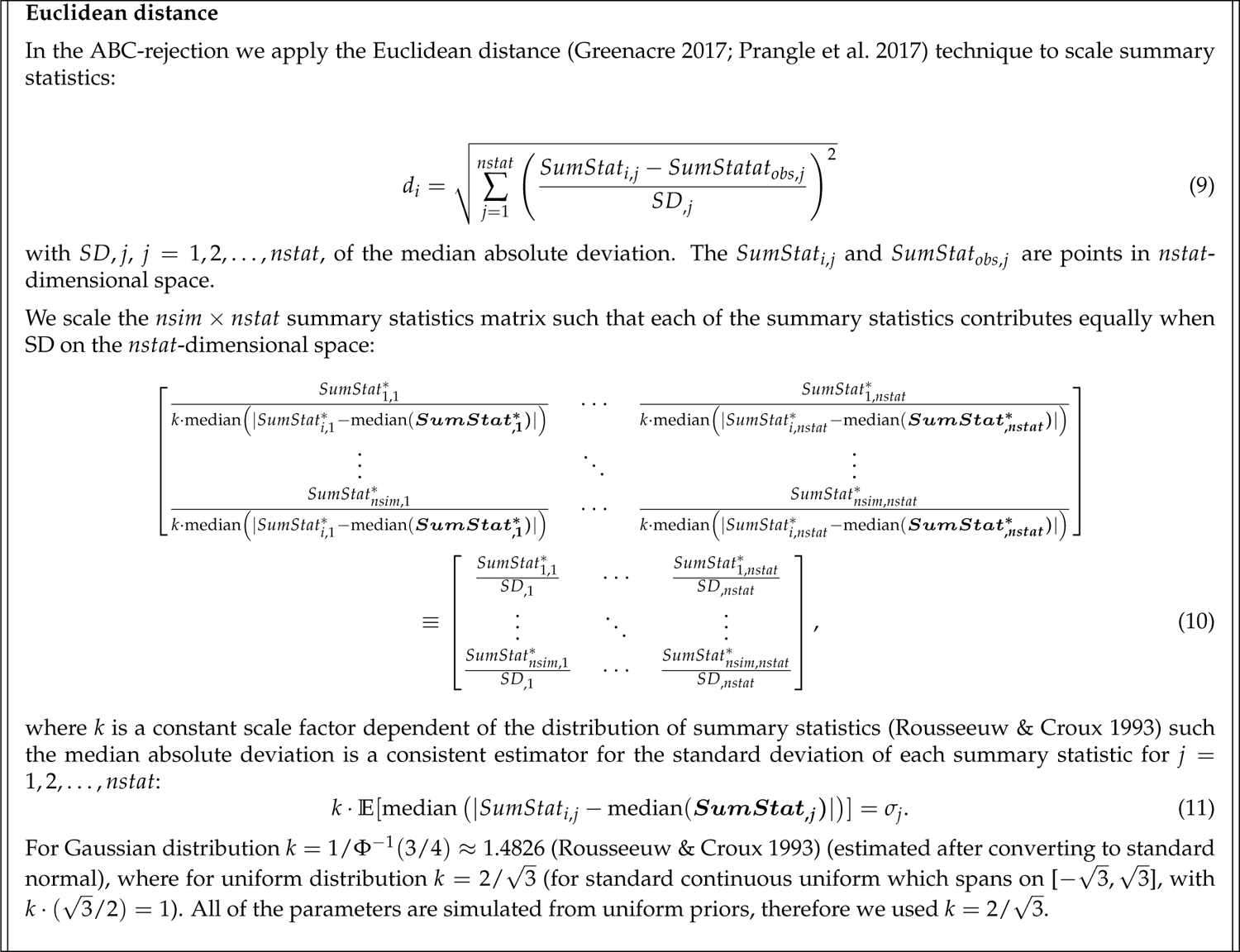
Euclidean distance equations.

**Table 5.**
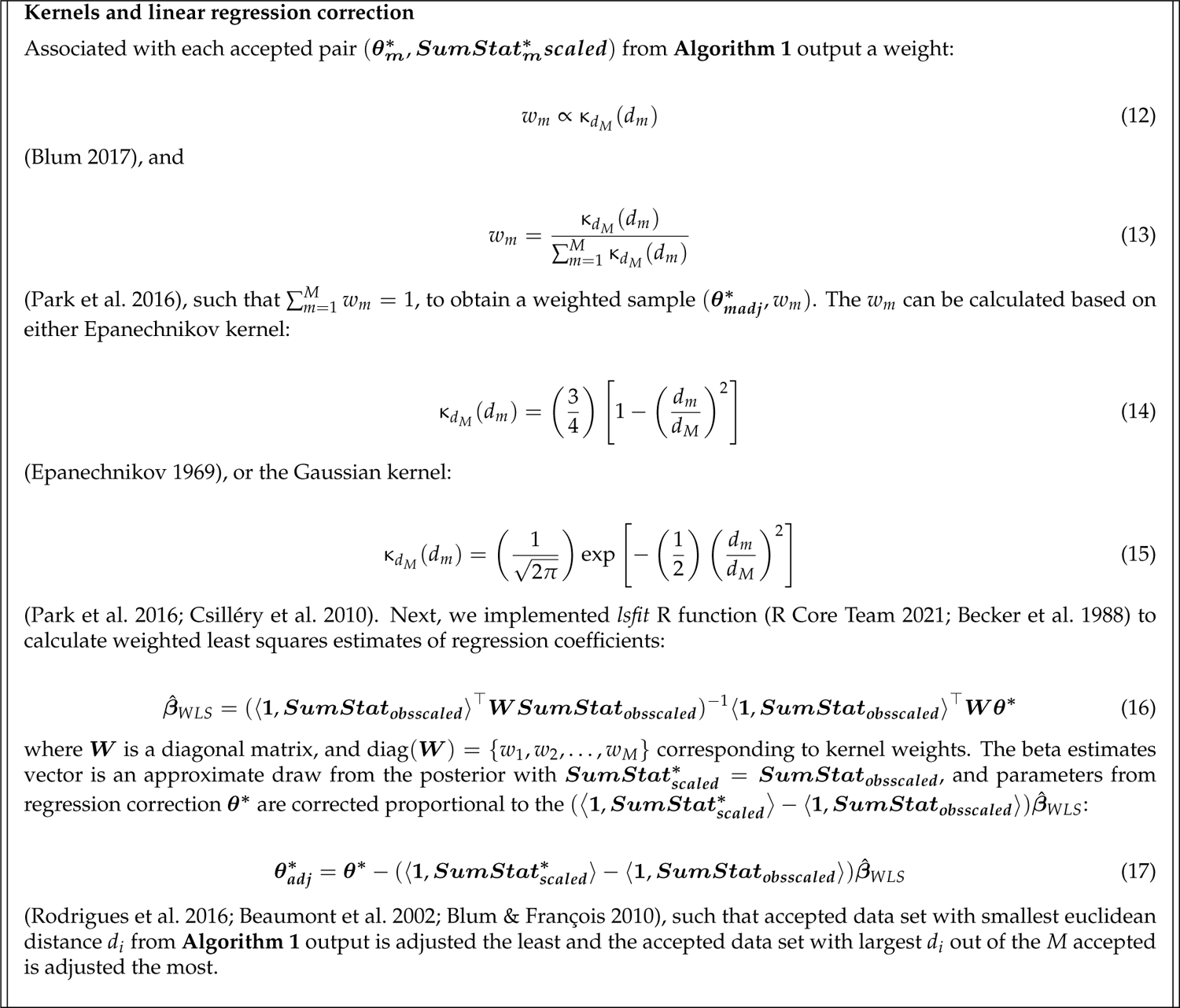
Kernel equations.

The experimental design part that differed from scenario 1 was: We performed 10, 000 ABC iterations instead of 100. Out of the potential 0, 1, or 2 loci under selection, the possible loci that the selection could act on were {*L*/3, *L*/2, 2*L*/3}, more specifically {33^rd^ SNP, 50^th^ SNP, 67^th^ SNP}.

For this small set of the only possible loci under selection, no outlier scan on observed summarized data was performed, which enabled us to reuse the same 100, 000 simulated data sets for the ABC iterations.

The experimental design of the parameters space for selection coefficients and rates of migration, with possible models of reproduction during migration generations of the model are seen in **Fig. 2**.

For the signature of the strength of selection due to mode of reproduction and strength of migration, we have evaluated *signF_ST_* summary statistic of each of 10, 000 observed data sets under four variable modes of reproduction (*sex*) and strength of migration (*m*) combinations: *sex* = 0 and *m* = 0, *sex* = 1 and *m* = 0, *sex* = 1 and *m* = 0.2, *sex* = 1 and *m* = 0.5, while equally probable seven combinations of 0,1,2 loci under selection within a set of {*L*/3, *L*/2, 2*L*/3} possible SNP loci (see **Fig. 2**). In **Fig. 3** we see a visible pattern in an increase in summary statistics values away from 0 (*signF_ST_* = 0 when the fixation index *F_ST_* = 0) at selected loci with a decrease in migration rate, and increase in genetic hitchhiking effect with an asexual mode of reproduction (*sex* = 0) given same *m* of *m* = 0.0.

For the signature of strength of selection due to a variable number of loci under selection, we have evaluated *signF_ST_* summary statistic of each of 10, 000 observed data sets where 0, 1, and 2 loci are under selection within a set of {*L*/3, *L*/2, 2*L*/3} possible SNP loci, while equally probable four combinations of mode of reproduction and migration described above (see **Fig. 2**). In **Fig. 4** we see a visible pattern in summary statistics values closest to 0 for no loci under selection, increase in the magnitude of summary statistics values away from 0 at single locus under selection, and in-between the magnitude of summary statistics values away from 0 and increase in genetic hitchhiking effect for two loci under selection.

In **Fig. 3** we see a break down of LD when recombination rate is present versus absent, given no migration rate, and in **Fig. 4** we see more break down in LD for one locus under selection instead of two loci, given same average genomic recombination rate, which is consistent with recombination as the primary source of LD break down (Qanbari 2020).

To further examine the signature of LD decay and distance between loci under selection, given fixed average genomic recombination rate, we looked at cases with loci under selection in closer proximity to each other. We evaluated effect of number of loci and distance between loci under selection for two loci cases, i.e. four combinations: none, one locus, two loci *L*/6 distance apart, and two loci *L*/3 distance apart, as well as each of seven loci combinations, i.e: none, *L*/3, *L*/2, 2*L*/3, (*L*/3, *L*/2), (*L*/3, 2*L*/3), (*L*/2, 2*L*/3) but we could not distinguish visually further differences than those seen in **Fig. 4** in observed summary statistics values due to position (seven combinations), nor the distance (four combinations) (see **Appendix C**).

In order to answer the question how well the signal we recover the signal of strength of selection, with variable selection coefficients priors, when the mode of reproduction, strength of migration, and number of loci under selection change, we evaluated the bias and the variability in the estimates of selection coefficients from the ABC. For this, we plotted the true observed selection coefficients versus: posterior medians, MSEs between true observed versus median of posteriors, variance of the posterior medians, and bias squared between true observed and posterior medians based on the ABC-linear regression Gaussian kernel. We picked Gaussian kernel because the PoPoolation software – a pipeline for analyzing pooled next generation sequencing data (Kofler et al. 2011) – uses Gaussian kernel smoothing (Hohenlohe et al. 2010). We assessed them based on four modes of reproduction and migration combinations, and based on number of loci under selection. For the mode of reproduction and migration (**Fig. 5**), we see a positive relationship between the magnitude of selection for strongest migration rate (*m* = 0.5) and variance as well bias (squared). The plot pattern for the number of loci under selection is more clear (**Fig. 6**), where we see a positive relationship between the magnitude of selection for two loci and variance as well bias (squared), and a change from positive to a negative relationship between the magnitude of selection for one locus and variance as well bias (squared) as the magnitude of selection increases. We also assessed Gaussian kernel performance with respect to Epanechnikov, ABC-rejection (**Algorithm 1**) versus Gaussian, and ABC-rejection (**Algorithm 1**) versus Epanechnikov. We compared to Epanechnikov kernel, and ABC-rejection and found no visual difference between kernel types, but an improvement in selection coefficient estimations for both Gaussian and Epanechnikov in comparison to ABC-rejection (**Appendix C**).

### 5.3 Observed Data, Model Simulations, and ABC for Potential loci under Selection Identified by Outlier Scan

In order to answer question how expansion of the parameter space for the positions of loci under selection contributes to accuracy in estimation of selection coefficients, we performed ABC evaluations for estimating strength of selection for model scenarios described in **Table 2**. The observed data sets had randomly selected 0, 1, or 2 loci under selection, and randomly selected loci under selection along *SNP* loci 1 to *L* (see **Model** section on parameter methodology). We identified candidates for loci under selection on the *n_ABC_* observed data sets via the outlier scan using the *F_ST_* summary statistic (see subsection 4.1) first, followed by the model simulations, then ABC.

We performed a total of four scenarios. For scenarios 1-3, we simulated same number of SNPs, average recombination rate, as described in method 1 in sub-subsection 5.2.1, and with same expected number of crossovers and average spacing of polymorphic sites on the genome equivalent to the biological yeast system as in scenario 4 (**Table 1**, and **Table 3** in **Appendix A**). Because of relatively low number of SNPs simulated, which referred which required lower computational time per simulated data set, we performed 100 ABC iterations (*n_ABC_*), which translated into 100 observed data sets (one ABC iteration per one observed data set), with *nsim* = 100, 000 unique data sets simulated conditional on outlier scan on potential locus under selection per each observed data set.

For scenario 1, the migration-selection cycle occurred every *t^∗^* = 5 generations, with total number of cycles *n_cycles_* = 4, and final generation *t _final_* = *n_cycles_t^∗^ −* 1 = 19. The prior parameter space of selection conditional on locus is *s_i_|I* = *i ∼* Unif[*−*0.25, 0.25]. This parameter space implies that if an overall population fitness *ω* is *ω* = 1 at *t* = 0, that is the average fitness of all carriers 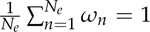 at *t* = 0, for extreme strength of selection if a carrier of one locus under selection has *s_i_|I* = *i* = *−*0.25 or *s_i_|I* = *i* = 0.25, it has 25% lower or higher fitness respectively than the population average at *t* = 0. For two loci under selection, this translates to 50% lower or higher fitness respectively than the population average at *t* = 0.

For scenario 2, the only parameters that differ from scenario 1 is the length of the selection cycle *t^∗^* = 50, a 10-fold increase from scenario 1, and the prior parameter space of selection conditional on locus is *s_i_|I* = *i ∼* Unif[*−*0.025, 0.025], a 10-fold decrease from scenario 1, as one would expect fixation to reach slower with weaker selection.

For scenario 3, the only parameter that differs from scenario 2 is number of cycles *n_cycles_* = 2, thus final generation *t _final_* = *n_cycles_t^∗^ −* 1 = 99. This scenario is to test whether the tested number of mixing (migration) times plays a significant role in the signature of selection in comparison to scenario 2.

For scenario 4, the recombination rate resembles closer to a genomic recombination rate of sequenced yeast data. The average recombination rate of *Saccharomyces cerevisiae* has been reported 3.5 *×* 10*^−^*^6^ Morgans/bp in the literature (Ruderfer et al. 2006), with inferred genome-wide recombination profiles from sequenced isolates from an advanced intercross line (AIL) to be as high as 3.0 *×* 10*^−^*^5^ Morgans/bp for a two-way cross at genome hotspots (Illingworth et al. 2013). Here, for the scenario resembling sequenced yeast, we fixed genome recombination rate of *r* = 2.0 *×* 10*^−^*^5^ Morgans/bp, a realistic value for recombination hotspots in yeast (Illingworth et al. 2013). The simulated number of SNPs for scenario 4 for the genomic chunk is *L* = 1, 500, around an average number of SNPs of a yeast cross for one-third a chromosome (Hether 2016), which still holds true that the expected number of cross-over events is *µ_nr_* = 4.95 (**Table 1**, and **Table 3** in **Appendix A**) as for scenarios 1-3, and as for the non-outlier scan earlier scenario with smaller parameter space of potential loci under selection ({*L*/3, *L*/2, 2*L*/3}). The pior parameter space of length of selection cycles, number of cycles (and thus final generation), and selection coefficients conditional on loci are equivalent to scenario 2 (*t^∗^* = 50, *n_cycles_* = 4, *t _final_* = *n_cycles_t^∗^ −* 1 = 199, *s_i_|I* = *i ∼* Unif[*−*0.025, 0.025]). Due to longer computational time because we simulated more SNPs in scenario 4 than in scenarios 1-3, we picked 10 instead of 100 data sets as observed and thus performed *n_ABC_* = 10 instead of *n_ABC_* = 100 ABC iterations.

Full list of scenarios with initial outlier scan for which the evaluations were performed are described in **Table 2**. The **Fig. 7** and **Fig. 8** show the summary statistics of observed data sets based on mode of reproduction and migration, and number of loci under selection respectively for each of the four scenarios. The ABC performance of MSE, variance and squared-bias between true selection parameter values under which the data are generated, and the values of posterior distributions for each of the four scenarios are shown in **Fig. 9**. In ABC analysis estimators are set to medians of the posterior distributions.

**Fig. 7.**
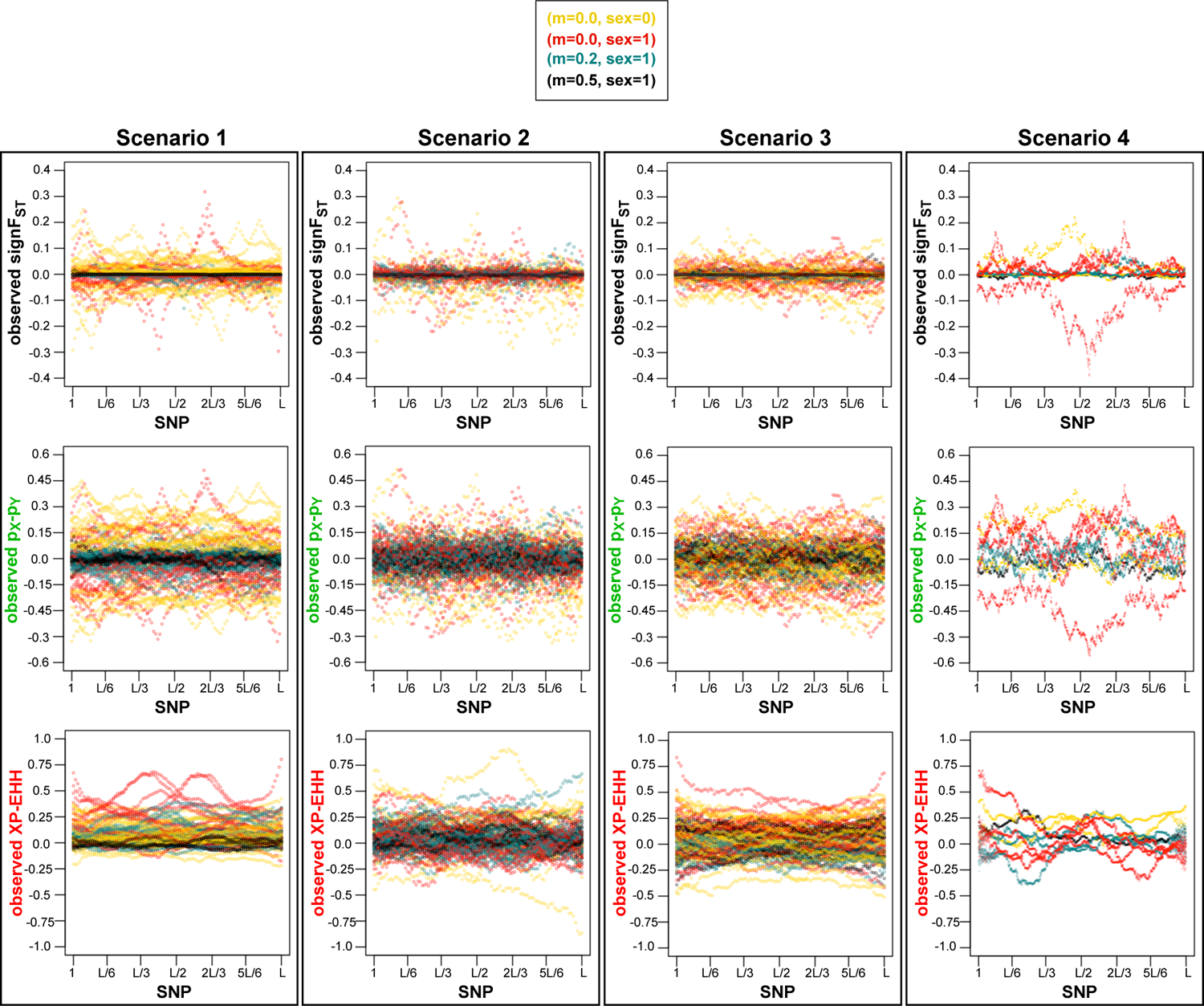
SNP loci vs. observed summary statistics for each of four scenarios with outlier scan method (Table 2) of simulator output data sets assumed as observed under four of migration (*m*) and mode of reproduction (*sex*) combinations selected randomly with equal probability: *sex* = 0 and *m* = 0.0 in yellow, *sex* = 1 and *m* = 0.0 in red, *sex* = 1 and *m* = 0.2 in turquoise, *sex* = 1 and *m* = 0.5 in black, and selected randomly with equal probability scenarios of loci under selection. Visible pattern in summary statistics values closest to 0 for strongest migration rate *m* = 0.5. Colors of observed summary statistics labels correspond to colors of plots in **Fig. 9**.

**Fig. 8.**
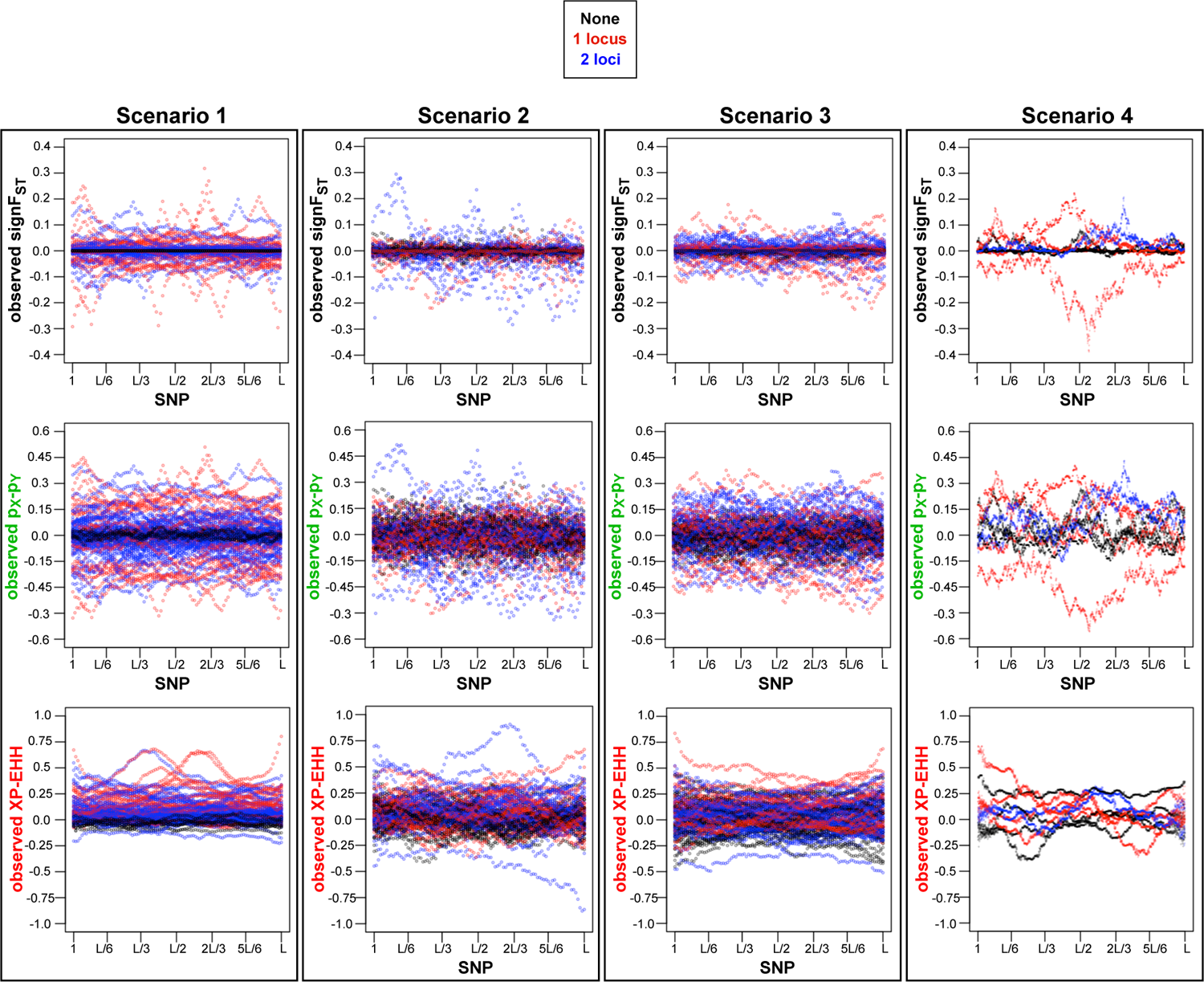
SNP loci vs. summary statistics for each of four scenarios with outlier scan method (Table 2) of simulator output data sets assumed as observed under four of migration (*m*) and mode of reproduction (*sex*) combinations selected randomly with equal probability scenarios of number loci under selection occuring within the loci identified by the outlier scan: none in black, one in red, and two blue. Visible pattern in summary statistics values closest to 0 for no loci under selection. Colors of observed summary statistics labels correspond to colors of plots in **Fig. 9**.

**Fig. 9.**
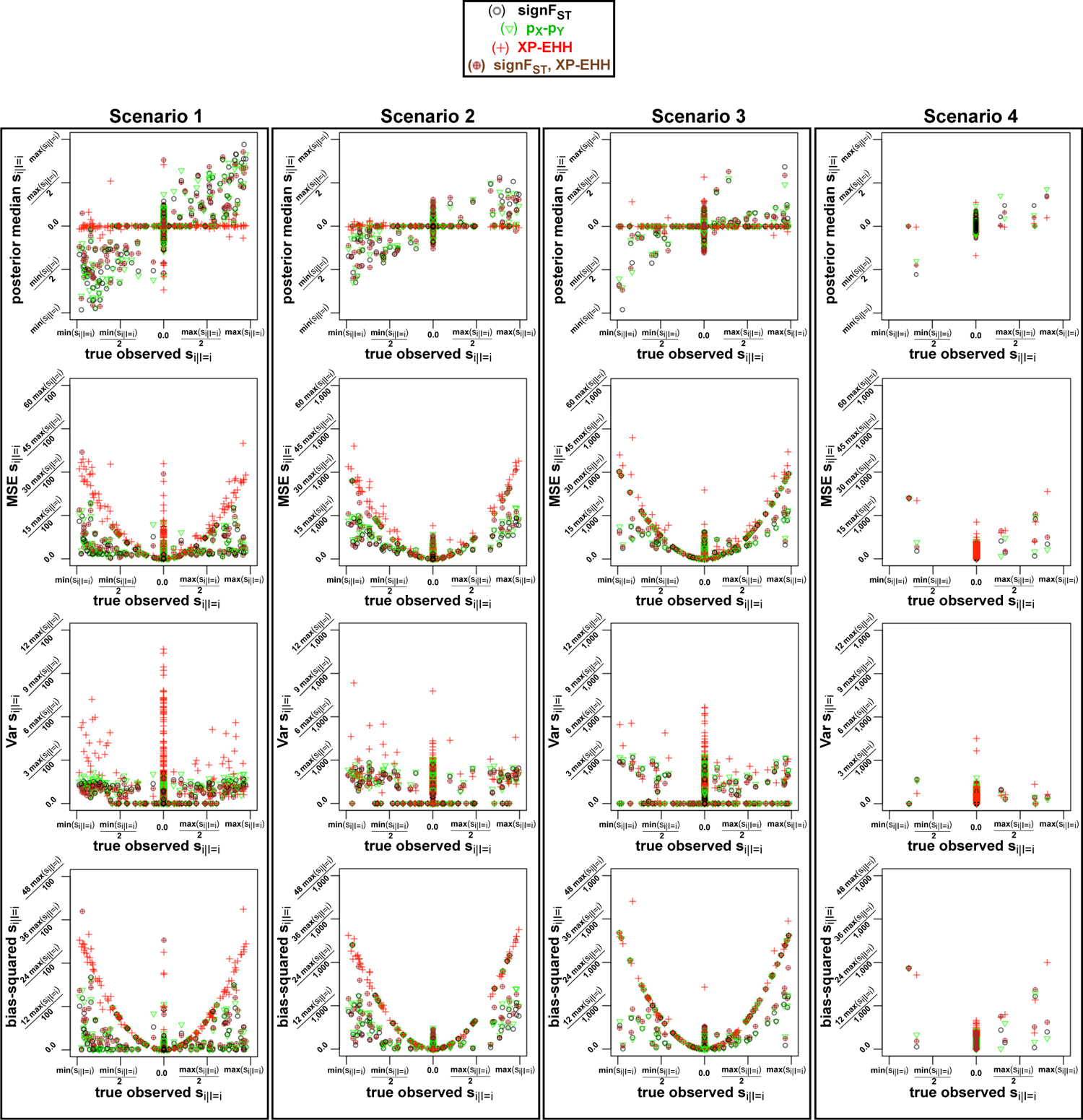
True parameter value under which the observed parameter is generated (x-axis) vs. median, MSE, variance, bias-squared for loci corresponding to SNP determined by initial outlier scan, followed by simulation of data sets and ABC from **Algorithm 2** with Gaussian kernel correction. Colors based on four combinations of summary statistics inputted to ABC are described in the figure legend. Plots for four scenarios, where in scenario 1 the range of selection coefficient is 10-fold larger ([0.25, 0.25]): thus the y-axis of the posterior median is 10-fold larger, the MSE, variance and bias-squared is (10-fold)-squared larger than for scenarios 2-4.

Out of the four scenarios, we see strongest pattern of observed summary statistics for scenario 1, where the range of selection coefficients is 10-fold greater, with shorter selection-migration cycles, characterized by sharp peaks (**Fig 7** and **Fig 8**). We also see that for scenario 1, the observed *XP − EHH* is almost exclusively positive. From expression 8 with population ***X*** in the numerator and population ***Y*** in the denominator, population ***X*** and ***Y*** are modeled under selection and under neutrality respectively. The numerator is larger due larger sum of the extended haplotype homozygosity decay around the locus the selection is acting on. In scenario 1 with stronger selection, and shorter selection-migration cycles, the genetic drift is expected to have smaller effect proportionally to strength of selection and length of selection-migration cycles.

For scenario 4 (resembling a biological yeast system), where 15-fold larger number of SNPs are simulated than for scenarios 1-3, we see peaks of observed summary statistics values with small stochastic effect between the SNPs unlike a more smooth pattern in observed summary statistics of neighboring SNPs in scenarios 1-3. This pattern resembles more so the pattern of the level of heterozygosity (Gallone et al. 2016) on domestication and divergence of *S. cerevisiae*, and less so in scenarios 1-3.

Besides the differences of patterns of observed summary statistics between the scenarios, we see a consistent pattern across the scenarios. The lowest level of divergence, expressed as the lowest magnitude of observed summary statistics, is seen for the strongest rate of migration (**Fig. 7**), and no loci under selection (**Fig. 8**).

After examining patterns of observed summary statistics by mode of reproduction with migration, and by number of loci under selection, we compared the ABC performance with these observed summary statistics by evaluating the variance of ABC posteriors, squared-bias between of the posterior estimates and true observed parameters. We find a clear pattern of posterior estimates to be less biased for scenario 1, where the range of selection coefficients is 10-fold of those in scenarios 2-4, and where the selection-migration cycles are shorter. Once explanation could be that short migration-selection cycles do not allow for the significant build-up of genetic drift, and/or that strong selection has too significant of effect for the genetic drift to act on the population.

## 6 Discussion

### 6.1 Application

By combining ABC methods that incorporate summary statistics on large simulation study from developed simulator, we present an approach of estimating model parameters, which can bypass evaluation of exact likelihood function. We show it on simulations with variable migration rates, modes of reproduction, and number of loci under selection, where fixation index summary statistics outperformed cross-population extended haplotype homozygosity in terms of precision and accuracy. We also recommend fixation index over the LD-based statistic cross-population extended haplotype homozygosity, because calculating the linkage disequilibria is computationally expensive (Alachiotis & Pavlidis 2016).

An important question about the application of these methods is for which part of the parameter space of the model the recovered signal about the selection coefficients well. In **Fig. 9** scenario 1 for instance the posterior median estimates of selection coefficients are closer to the values of the true observed selection coefficients in compare to scenarios 2-4. In scenario 1, the migration- selection cycles are 10-fold smaller, and the selection coefficients range is 10-fold larger thus affected less by the genetic drift, supporting that selection can only be assessed if it is high enough to outperform the effect of genetic drift (Mariac et al. 2016). The shorter migration-selection cycles with overall less generations simulated, took less time to simulate. We showed in scenario 4 that our model is scalable to recombination rate parameter resembling sequenced yeast data (Illingworth et al. 2013), with same expected number of recombination events (**Table 1**, **Table 3** in **Appendix A**) as for scenarios 1-3. If the parameter space of possible loci under selection can be assumed to those in **Fig. 2**, a more robust number of ABC iterations on data resembling biological yeast recombination rate is feasible in terms of computational time.

### 6.2 Recombination Rate

A limitation of the study where the main focus was estimation of selection coefficients, was fixing the recombination rate to an expacted number of 4.95 recombination events (**Table 1**, **Table 3** in **Appendix A**) for recombination rates 3.0 *×* 10*^−^*^4^ Morgans/bp and resembling biological data 2.0 *×* 10*^−^*^5^ Morgans/bp (Illingworth et al. 2013) for 100 and 1,500 SNPs respectively. We fixed the recombination rate to same for all simulations, such that the parameter *r* represents the average recombination rate on the simulated genome section. We explain out motive for fixing the recombination rate below.

With our model we attempted to estimate strength of selection with variable recombination rates of simulated data sets, however, we were unable to get consistent estimates. While the buildup of LD (i.e. the correlation between nearby variants of alleles as opposed to random association of alleles (Slatkin 2008)) can be a result of several population genetic forces, recombination is the only primary method to break it down (Qanbari 2020). The absence of recombination between sites under selection will reduce the overall effectiveness of selection, a phenomenon known as the Hill-Robertson (HR) effect (Hill & Robertson 1966; Comeron et al. 2008; Qanbari 2020). Our main focus was estimation of strength of selection, with successfully applied variability in mode of reproduction, and in strength of migration, but not with variability in recombination rate. As the *XP − EHH* was least effective in estimation of selection with fixed recombination, and *XP − EHH* has been used to measure the haplotype lengths between two populations (Sabeti et al. 2002; Weigand & Leese 2018), we would expect *XP − EHH* to perform better in estimation of recombination without taking into account varying strength of selection, mode of reproduction and migration combined.

Future work direction would be an exploration of variable recombination rates.

## Appendix A

### Algorithm 3

Algorithm for population process given in **Fig. 1**. Input: Effective population sizes *N_e_*, for ***X*** and ***Y*** ; Genome size *L*, for each individual; Vector of loci of selected sites, ***L_s_***; Vector of alleles at selected sites, *a*; recombination rate of parental genomes, *r*; strength of selection per locus *s_i_*; migration rate from population ***X*** to population ***Y***, *m_XY_*; migration rate from population *Y* to population ***X***, *m_YX_*; migration event, between *t^∗^ −* 1 and *t^∗^*, SNP spacing on the genome, *SNP_spacing_* (assumed equally spaced SNPs). Parts I, II, A, B and C of **Fig. 1** are performed on 2–5, 6–20, 21–30, 31–45, and 46–48 respectively.

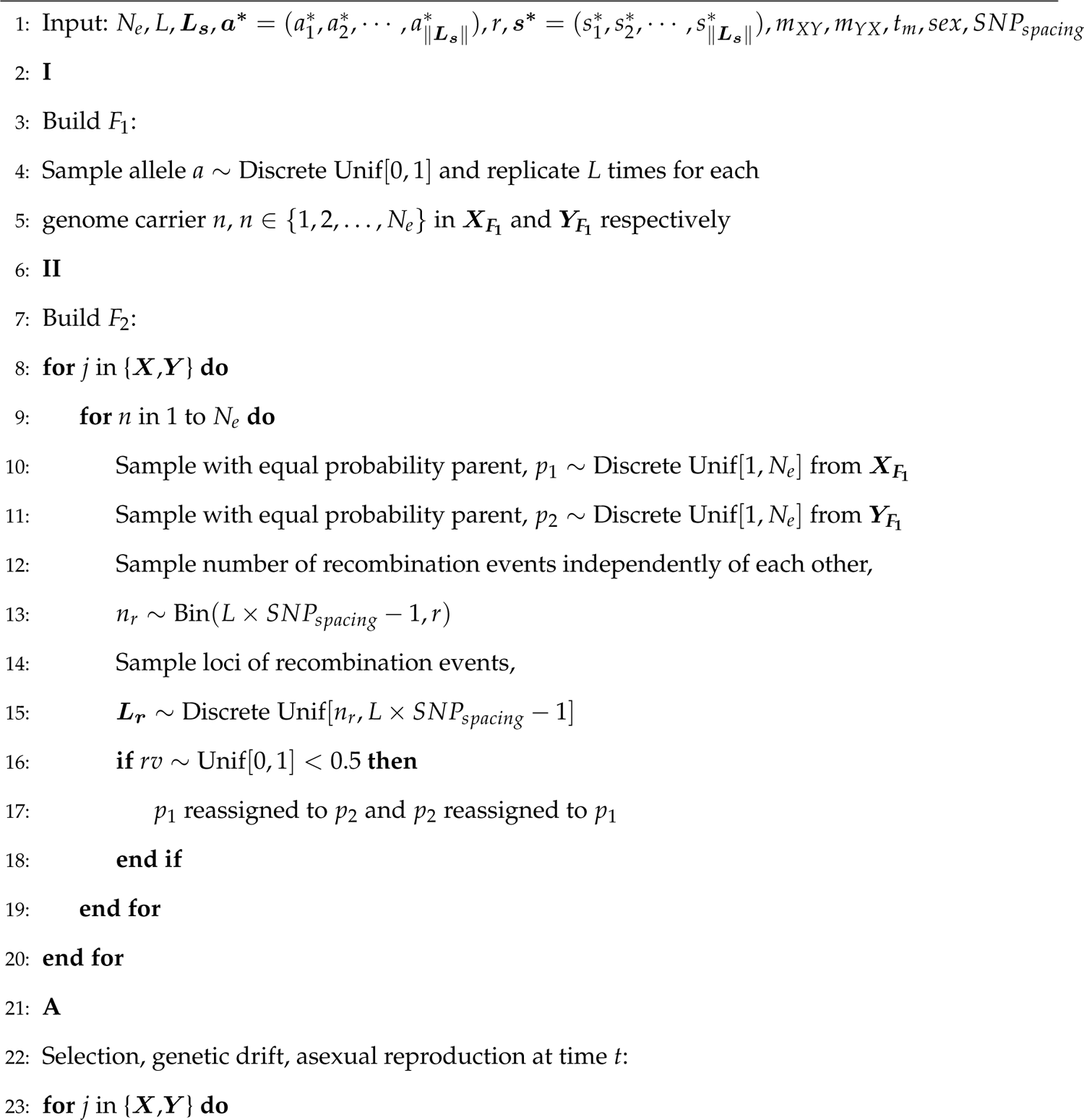
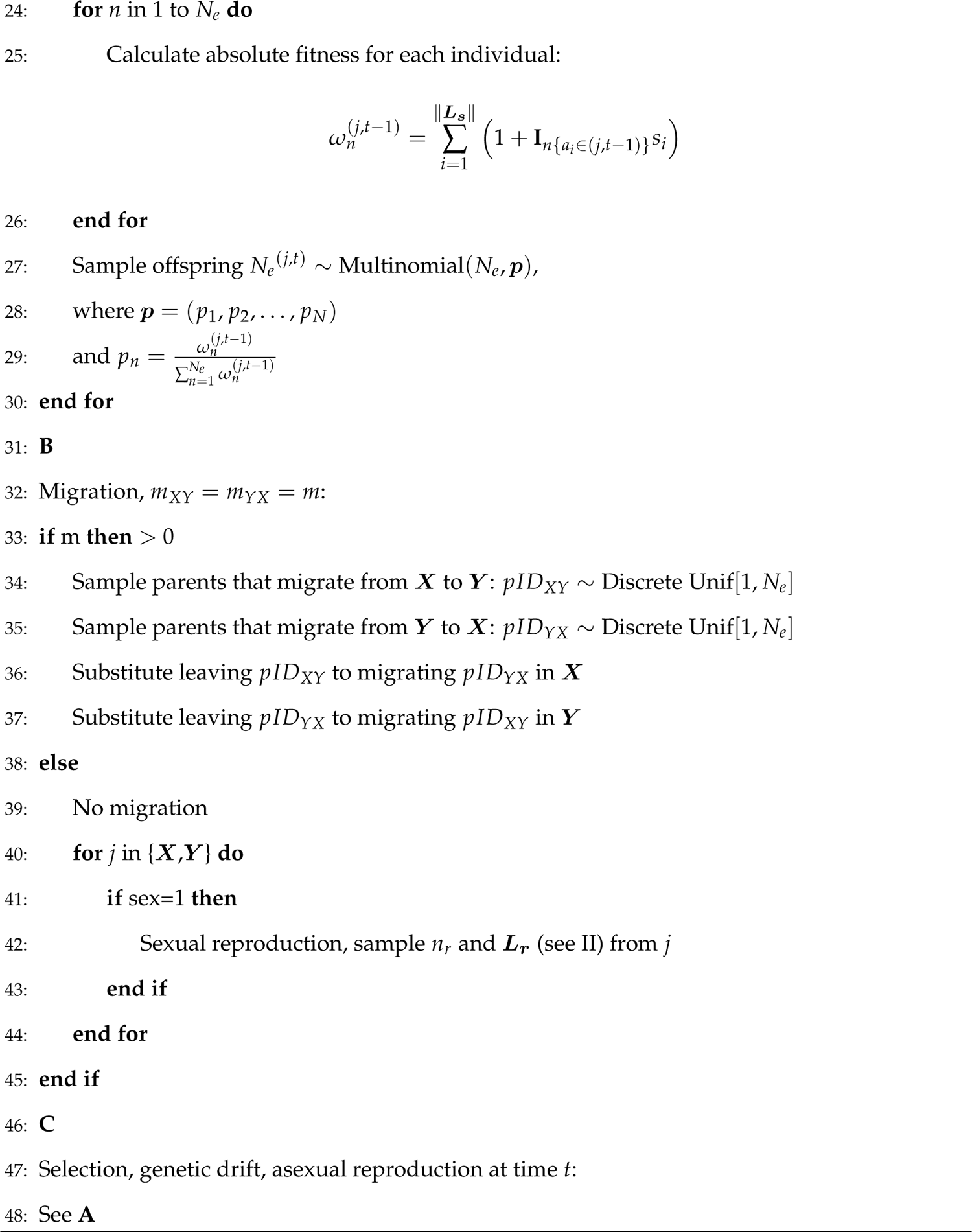

### Algorithm 4

ABC-rejection algorithm for summary statistics calculated from datasets simulated from the **Simulator** and their corresponding parameters from the prior distribution. Input: proportion of simulations to accept *tol*, number of simulated data sets *nsim*, number of summary statistics per simulation *nstat*.

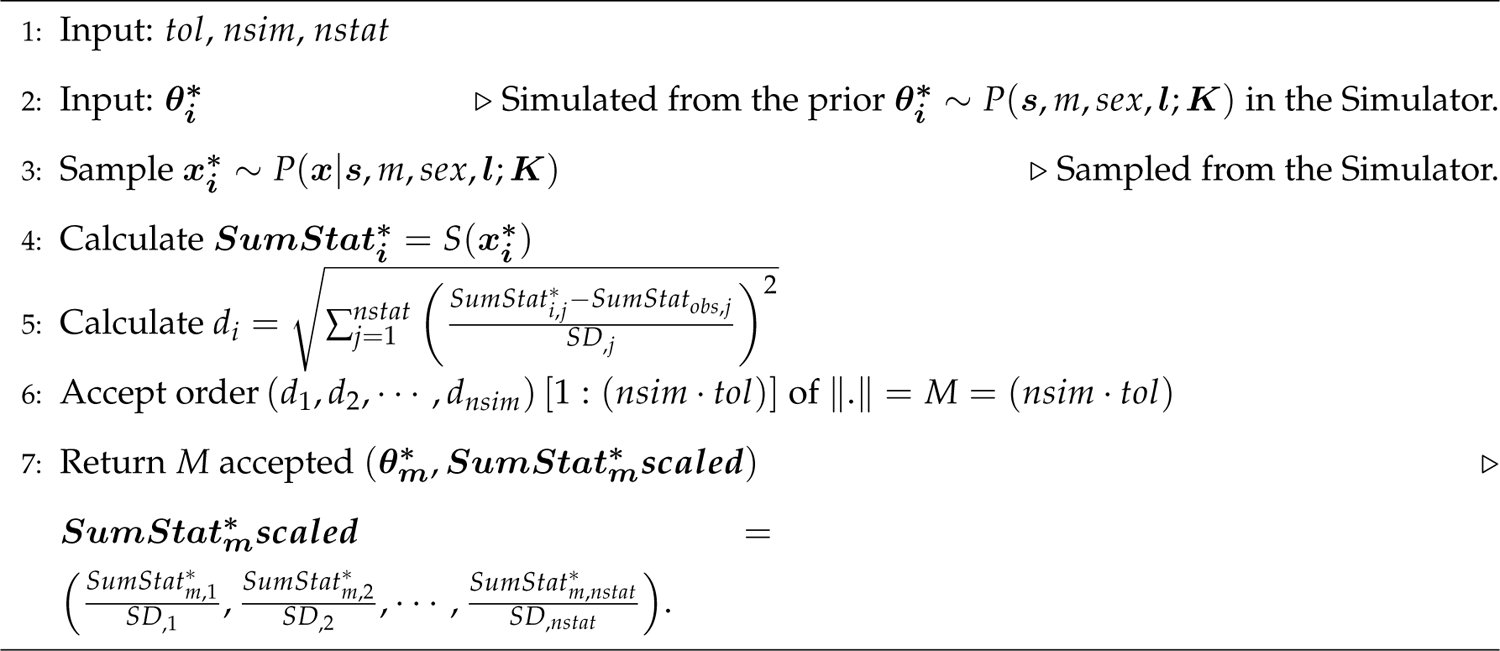

### Algorithm 5

ABC-linear regression correction algorithm performed on the output of the ABC-rejection. Input: ***SumStat_obsscaled_***,, *M* accepted ***SumStat*scaled***, *M* accepted ***θ^∗^*** from ABC-rejection.

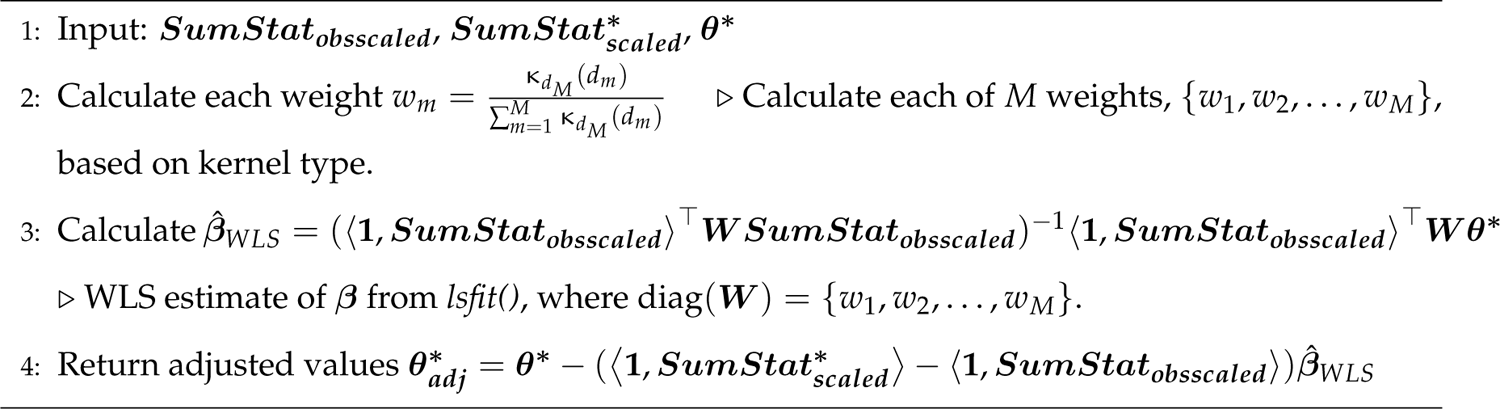

## Appendix B

Detailed description of the assessment of genetic drift on the model from subsection 4.1.

In order to assess the effect of genetic drift and most informative sample summary statistic, we did the following in order to compare mean square errors (MSEs) of *s*: (1) we examined change in difference of allele frequency between population ***X*** and ***Y*** for every generation of the model for a single locus under selection (midpoint SNP), with reduced the model to *L* = 20 SNPs, with selection at locus at *L*/2^th^ SNP rounded up of *s_L_*_/2_ = 0.1, migration *m* = 0.1, spacing of SNPS every 165 loci (*SNP_spacing_* = 165), and recombination rate *r* = 2.880 *×* 10*^−^*^4^; (2) we compared the stochasticity of a single randomly chosen simulation to an average of the change in allele frequencies per SNP from 100,000 simulations; (3) we simulated a single autosomal locus (a deterministic case, infinite population size) (Gillespie 2004) under same strength of selection, with equal proportions of allele 0 and 1 (expected values from the simulated model) – both 0.5 – in both populations at generation *t* = 0 (equivalent to *t* = 0 in **Part A** in **Fig. 1**), and we updated frequencies per generation to:

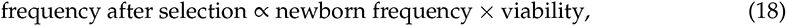

We show negligent effect of genetic drift and effective population size of simulations on the difference in allele frequency *p* between population ***X*** and population ***Y***, (*p_X_ − p_Y_*), summary statistics at locus under selection when we average our summary statistics at each SNP over 100,000 simulations. By removing the stochastic effect we can deterministically identify most informative summary statistic via ABC approach. Here we track (*p_X_ − p_Y_*) per generation based on the model. **Fig. 10** (main, left plot) shows mean of (*p_X_ − p_Y_*) summary statistics of 100,000 simulations tracked from *t* = 0 to generation at end of the simulation. There, we see linkage disequilibrium between locus under selection (red line) and the neighboring loci (black lines), proportional to the distance between locus under selection and those that are not. The three drops in (*p_X_ − p_Y_*) summary statistics values correspond to the migration generations where the populations mixing occurs and therefore less difference seen in the proportion of alleles between populations ***X*** and ***Y*** . In the upper right plot we superimposed the red plot from the main, left plot and deterministically derived (*p_X_ − p_Y_*) for a single locus case (Gillespie 2004) for haploid case with applied migration every 50^th^ generation in green. We see that we cannot distinguish between single locus deterministically derived (*p_X_ − p_Y_*) values, and the red line (locus under selection, main, left plot) of the mean of (*p_X_ − p_Y_*) summary statistics of 100,000 simulations. For reference how stochastic a single simulation can be, we randomly picked one of the 100,000 simulations and plotted the (*p_X_ − p_Y_*) values (lower right plot). This visual representation verifies that with very large number of simulations the values of summary statistics (*p_X_ − p_Y_*) converge to deterministic (expected) values, and the signal of selection has an effect on the nearby loci (linkage disequilibrium effect).

**Fig. 10.**
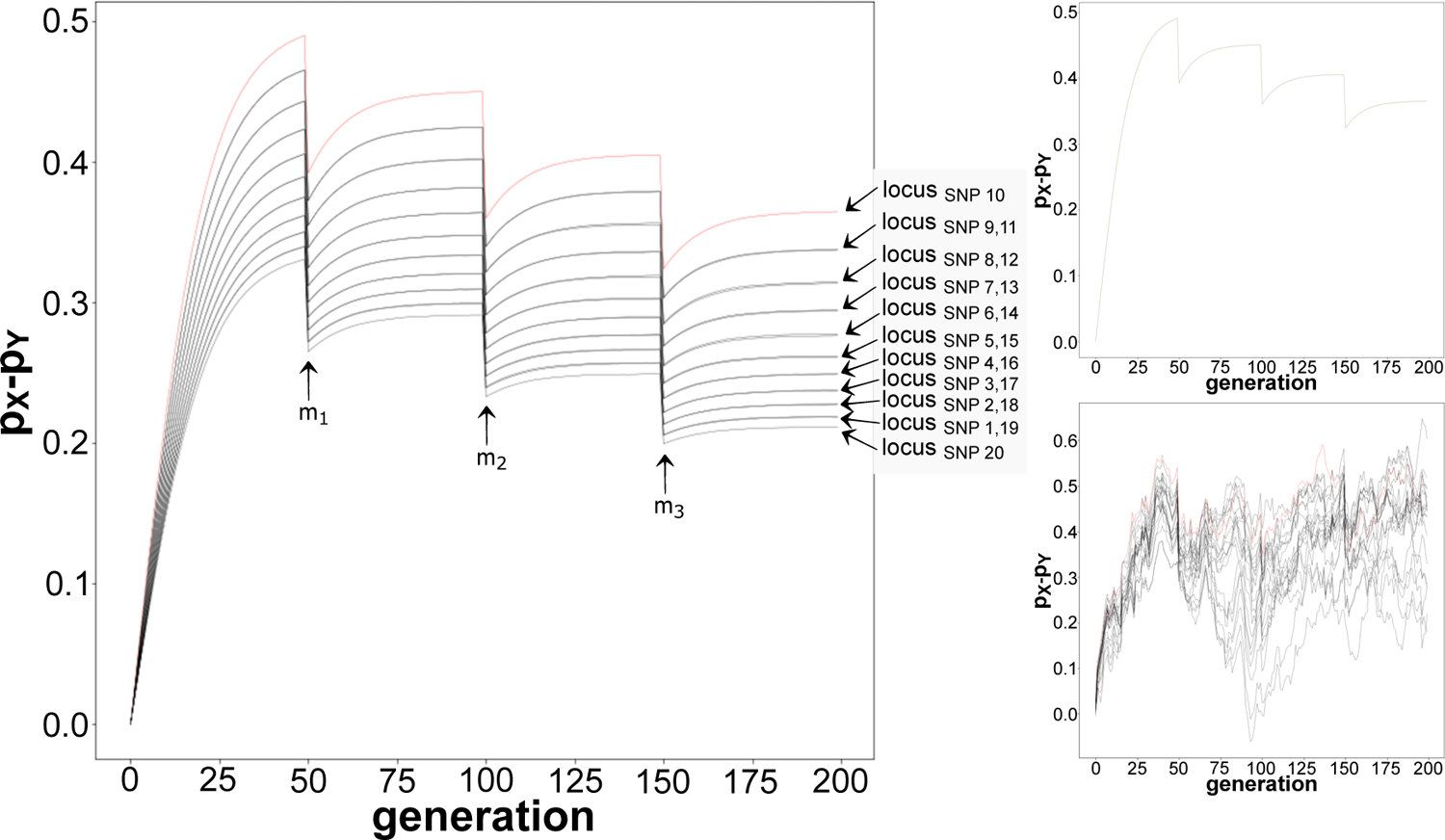
Mean of statistics (*p_X_ p_Y_*) per locus at SNP 1 to *L* = 20 for *nsim* = 100, 000 simulated data sets, with selection at locus at SNP 10, decrease in (*p_X_ p_Y_*) proportional to distance away from locus under selection, and with drops of (*p_X_ p_Y_*) values at migration *m* generations (left). Superimposed expected values and mean of 100, 000 simulated data sets of (*p_X_ p_Y_*) at locus under selection (upper right). Visible genetic drift for a randomly chosen simulated data set.

## Appendix C

Supplemental figures.

**Fig. 11.**
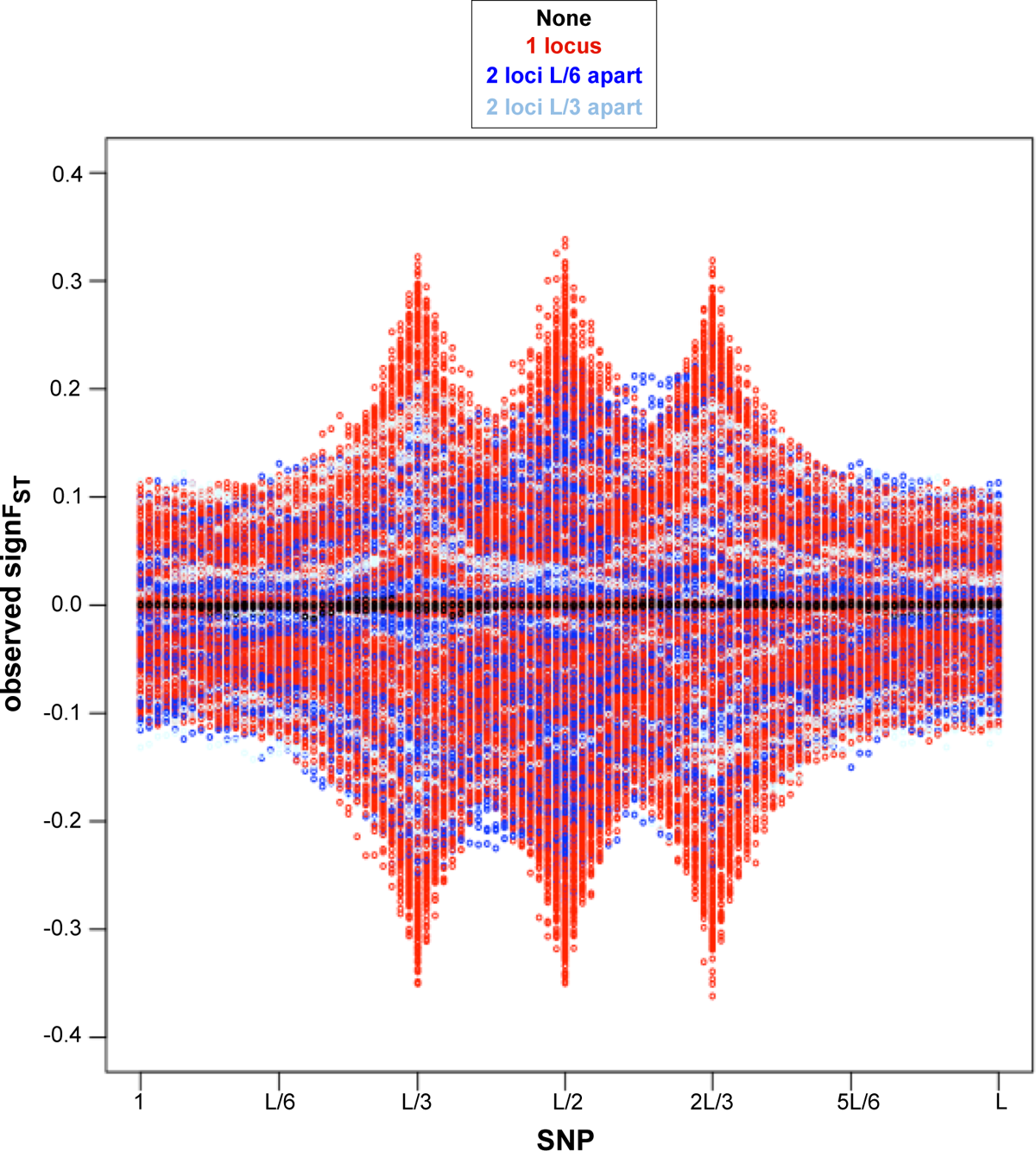
SNP loci vs. *signF_ST_* summary statistics for each of 10, 000 simulator output data sets assumed as observed under four of migration (*m*) and mode of reproduction (*sex*) combinations selected randomly with equal probability scenarios of loci under selection: none in black, one locus (*L*/3, or *L*/2, or 2*L*/3) in red, two loci *L*/6 distance apart in navy blue (*L*/3 with *L*/2, or *L*/2 with 2*L*/3), two loci *L*/6 distance apart in sky blue (*L*/3 with 2*L*/3), as seen in **Fig. 2**. Visible pattern in summary statistics values closest to 0 for no loci under selection, increase in summary statistics values away from 0 at single locus under selection, and in-between the magnitude of summary statistics values away from 0 and increase in genetic hitchhiking effect for two loci under selection but unclear of the observed summary statistic pattern difference between two loci *L*/6 (navy blue) versus *L*/3 (sky blue) distance apart.

**Fig. 12.**
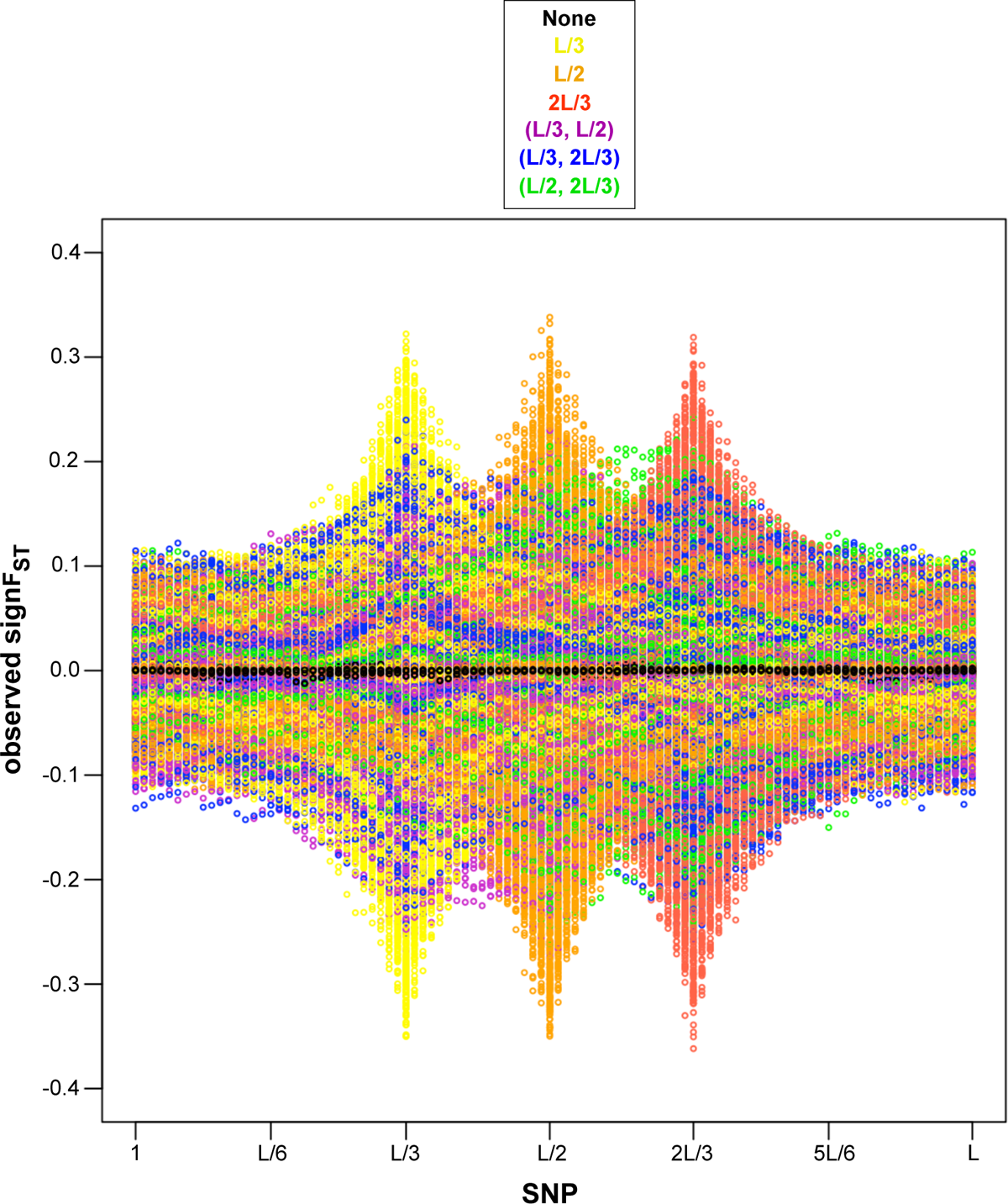
SNP loci vs. *signF_ST_* summary statistics for each of 10, 000 simulator output data sets assumed as observed under four of migration (*m*) and mode of reproduction (*sex*) combinations selected randomly with equal probability scenarios of loci under selection: none in black, *L*/3 in yellow, *L*/2 in orange, 2*L*/3 in red, *L*/3 with *L*/2 in purple, *L*/3 with 2*L*/3 in navy blue, and *L*/2 with 2*l*/3 in green, as seen in **Fig. 2**. Visible pattern in summary statistics values closest to 0 for no loci under selection, an increase in summary statistics values away from 0 at single locus under selection, and in-between the magnitude of summary statistics values away from 0 and increase in genetic hitchhiking effect for two loci under selection but unclear of the observed summary statistic pattern difference between two loci combinations of *L*/3 with *L*/2 (purple), versus *L*/3 with 2*L*/3 (navy blue), versus *L*/2 with 2*L*/3 (green).

**Fig. 13.**
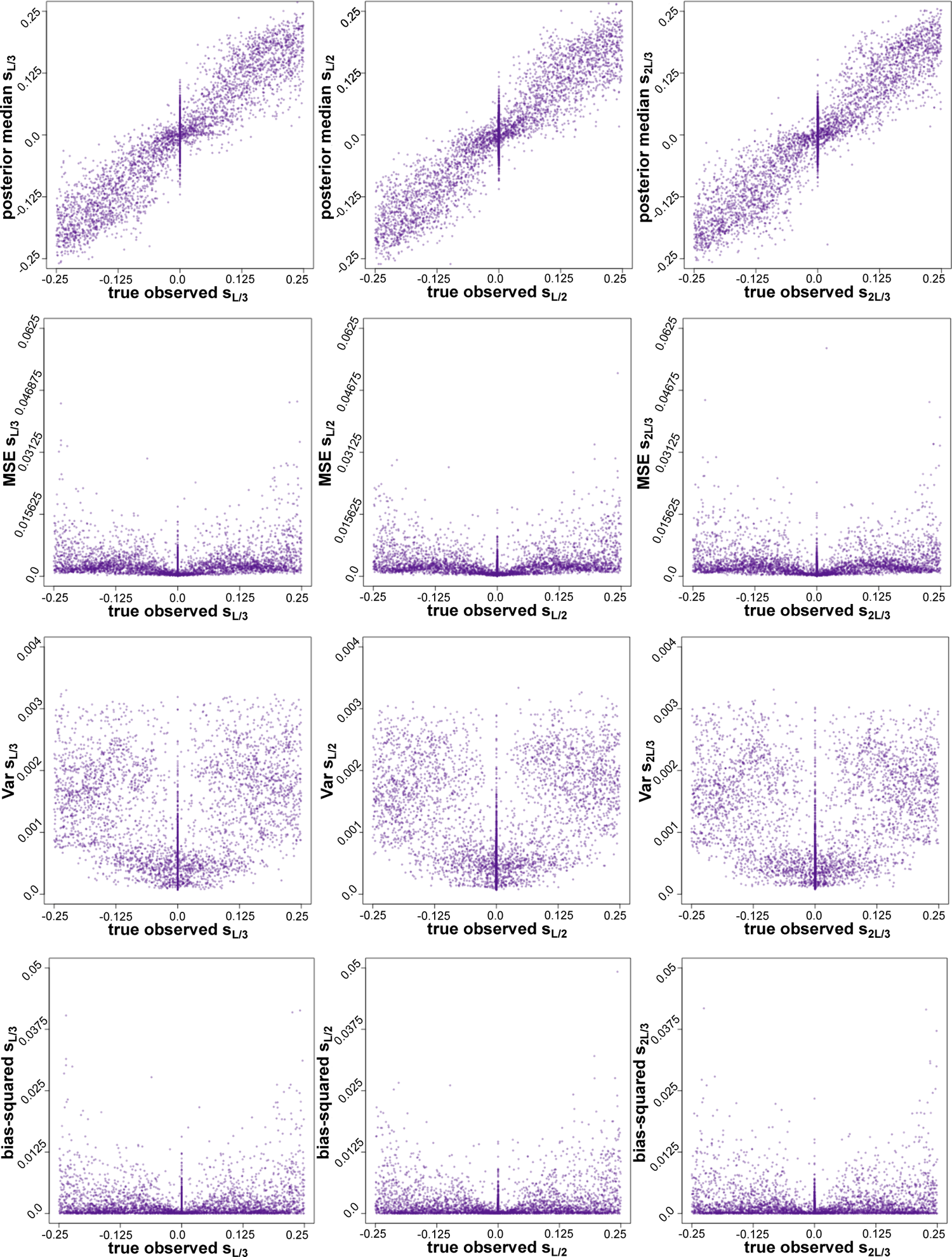
True parameter value under which the observed parameter is generated (x-axis) vs. median, MSE, variance, bias-squared for (*L*/3)^th^, (*L*/2)^th^ and (2^5^*L*^3^/3)^th^ SNP respectively from 10,000 ABC tests from *signF_ST_* summary statistics from **Algorithm 2** with Gaussian kernel.

**Fig. 14.**
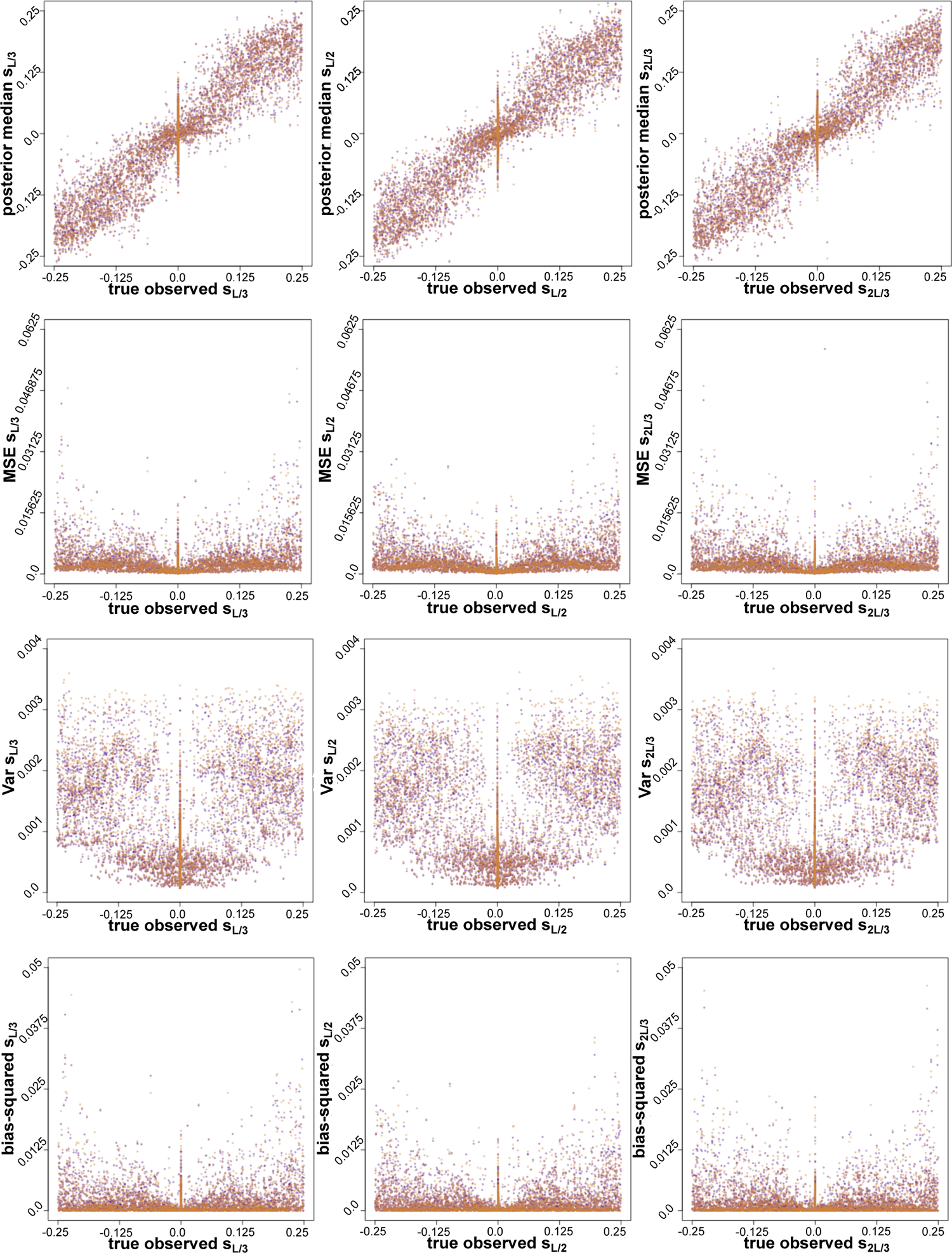
True parameter value under which the observed parameter is generated (x-axis) vs. median, MSE, variance, bias-squared for (*L*/3)^th^, (*L*/2)^th^ and (2^5^*L*^4^/3)^th^ SNP respectively from 10,000 ABC tests from *signF_ST_* summary statistics from **Algorithm 2** with Gaussian kernel (purple) and Epanechnikov kernel (orange).

**Fig. 15.**
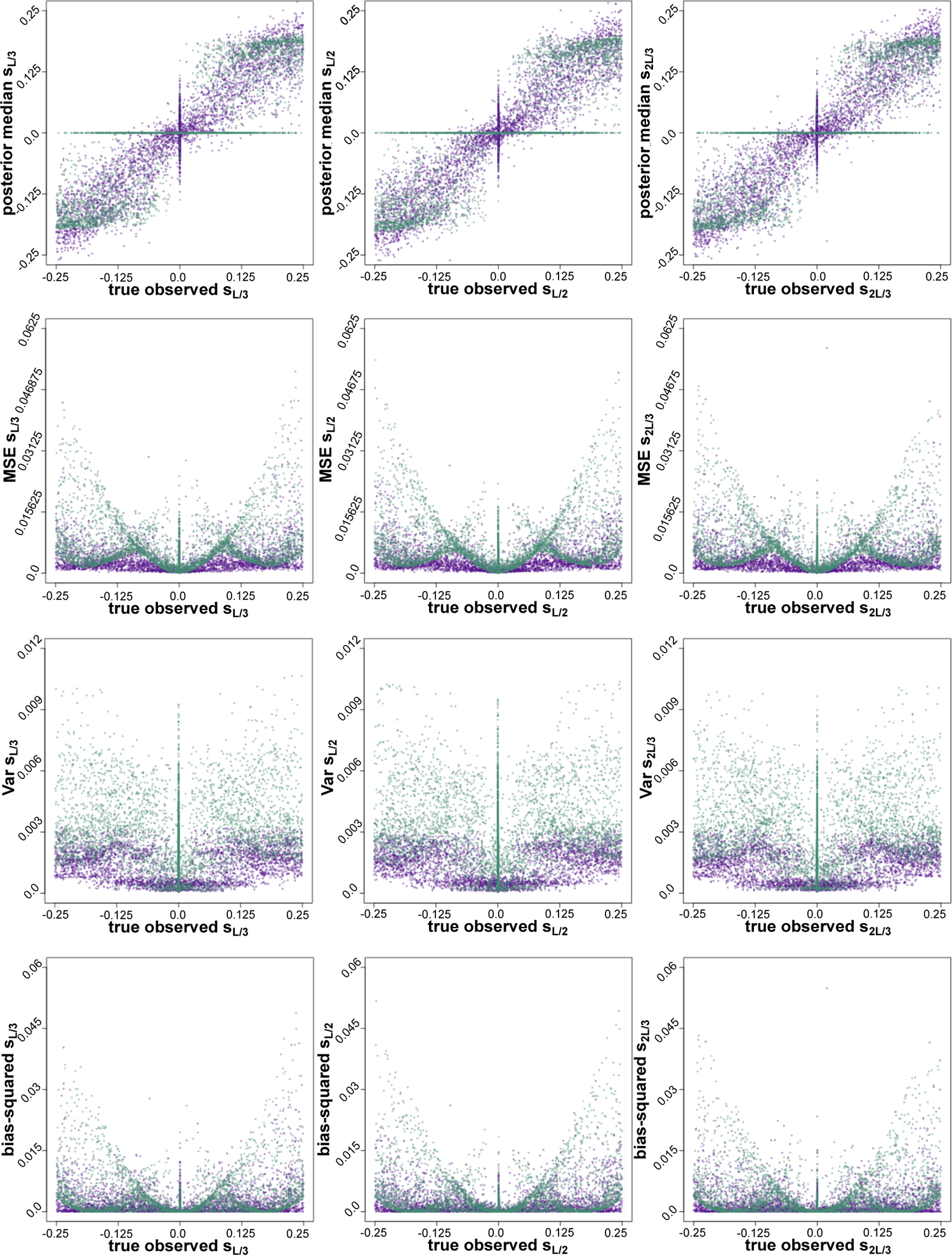
True parameter value under which the observed parameter is generated (x-axis) vs. median, MSE, variance, bias-squared for (*L*/3)^th^, (*L*/2)^th^ and (2^5^*L*^5^/3)^th^ SNP respectively from 10,000 ABC tests from *signF_ST_* summary statistics from **Algorithm 2** with Gaussian kernel (purple) and from **Algorithm 1** (green).

**Fig. 16.**
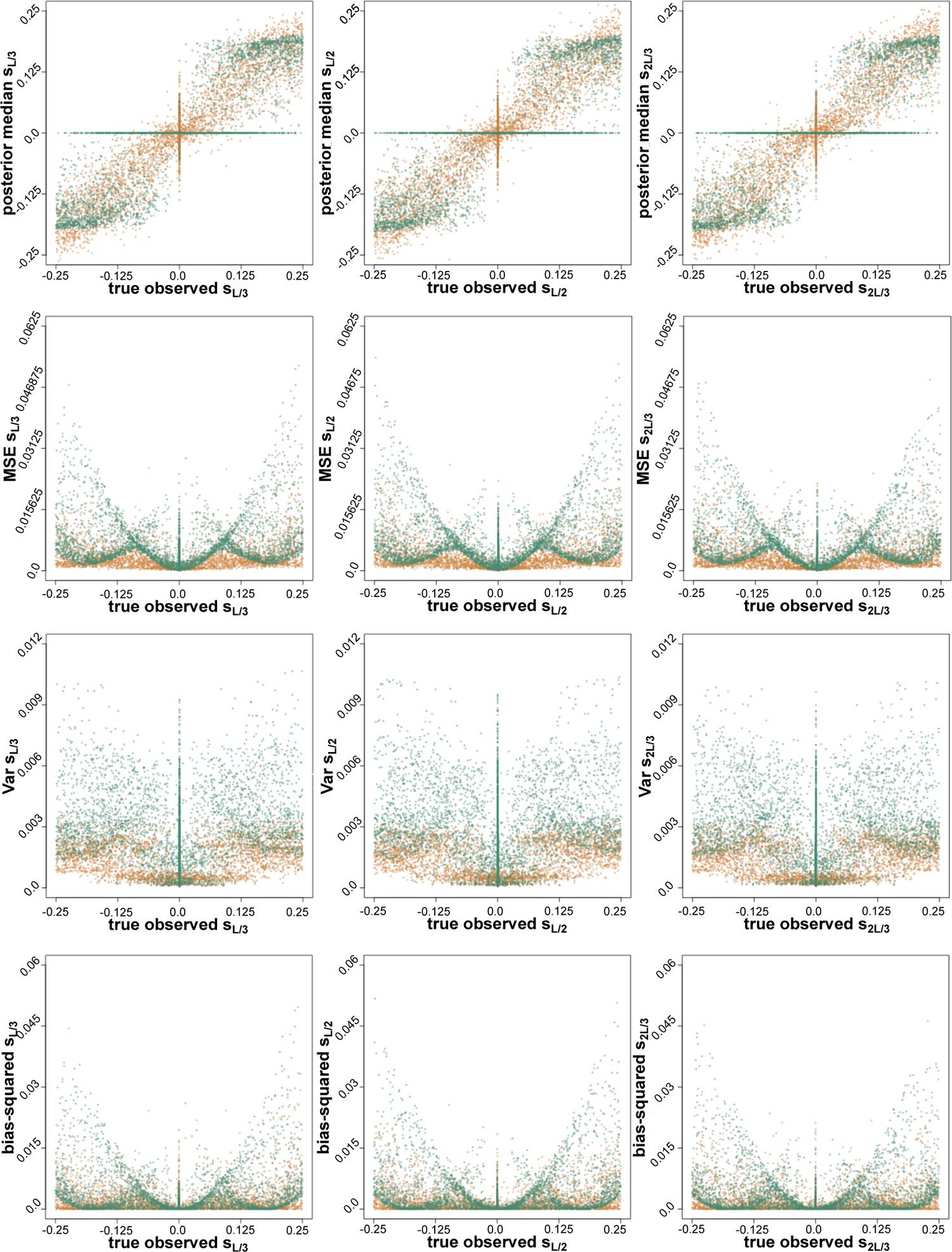
True parameter value under which the observed parameter is generated (x-axis) vs. median, MSE, variance, bias-squared for (*L*/3)^th^, (*L*/2)^th^ and (2^5^*L*^6^/3)^th^ SNP respectively from 10,000 ABC tests from *signF_ST_* summary statistics from **Algorithm 2** with Epanechnikov kernel (orange) and from **Algorithm 1** (green).

